# Sleep reactivation of positive self-evaluation reveals neural-evaluative alignment that varies with depressive symptoms

**DOI:** 10.64898/2026.07.26.740831

**Authors:** Ziqing Yao, Danni Chen, Jing Liu, Jinwen Wei, Tao Xia, Xibo Zuo, Chris Xie Chen, Rachel Ngan Yin Chan, Shirley Xin Li, Pengmin Qin, Tifei Yuan, Zhe Zhang, Tatia M.C. Lee, Yina Ma, Xiaoqing Hu

**Affiliations:** Department of Psychology, The University of Hong Kong, Hong Kong SAR, China; School of Psychology, South China Normal University, Guangzhou, China; Department of Physics, Hong Kong Baptist University, Kowloon Tong, Hong Kong, China; Institute of Psychology Chinese Academy of Sciences, Beijing, China; Li Chiu Kong Family Sleep Assessment Unit, Department of Psychiatry, Faculty of Medicine, The Chinese University of Hong Kong, Hong Kong SAR, China; Shanghai Key Laboratory of Psychotic Disorders, Brain Health Institute, National Center for Mental Disorders, Shanghai Mental Health Center, Shanghai Jiao Tong University School of Medicine and School of Psychology, Shanghai 200030, China; Center for Excellence in Brain Science and Intelligence Technology, Chinese Academy of Sciences; Laboratory of Neuropsychology and Human Neuroscience, Department of Psychology, The University of Hong Kong, Hong Kong SAR, China; Faculty of Psychology, IDG/McGovern Institute for Brain Research, Beijing Normal University, Beijing, China; Beijing Institute for Brain Research, Beijing, China; The HKU-Shenzhen Institute of Research and Innovation, Shenzhen, China

**Author notes:** Correspondences: Yina Ma, Xiaoqing Hu.

**Keywords:** targeted memory reactivation, positive self-evaluation, NREM sleep, depressive symptoms, representational similarity analysis

## Abstract

Positive self-evaluation underpins resilience and mental health, yet how to enhance positive self-evaluation and the underlying neurocognitive mechanisms remain poorly understood. Existing data mostly examined self-evaluative processing of the wakeful minds, with the role of sleep remain unexplored. Here, we demonstrated that reactivation of positive self-evaluation during sleep revealed fine-grained neural-evaluative alignment that varies with depressive symptoms.

We found that targeted reactivation of positive personality traits during non-rapid eye movement (NREM) sleep enhanced positive self-evaluation, but this benefit diminished with elevated depressive symptoms. Moreover, cueing positive traits during NREM sleep elicited trait-specific, 11-16 Hz EEG sigma power-based neural representations that aligned with post-sleep positive self-evaluative representational structures, as revealed by representational similarity analyses.

Notably, individuals with elevated depressive symptoms exhibited weaker neural-evaluative representational alignments, identifying a maladaptive self-evaluative processing that bridges across sleep and wakefulness states. These findings provide mechanistic insights on how fine-grained neural-evaluative alignment can substantiate positive self-evaluation, and why it was disrupted as depressive symptoms increased.

**Highlights:**

- Positive-trait reactivation during NREM sleep enhances positive self-evaluation
- This behavioral benefit decreases as depressive symptoms increase
- Sleep cueing evokes trait-specific sigma-band neural representations
- Neural–evaluative alignment weakens as depressive symptoms increase

## Introduction

Positive self-evaluation, the tendency to view one’s traits, abilities and social worth in a favourable light, is a cornerstone underpinning psychological well-being, resilience, and social functioning (Symons & Johnson, 1997; Cohen & Sherman, 2014; Howell, 2017; Sowislo & Orth, 2013; Orth & Robins, 2022). By contrast, diminished positive self-evaluation lead to self-doubts and senses of worthlessness, contributing to depression (Collins & Winer, 2024; Disner et al., 2011; Dozois & Beck, 2008; Weisenburger et al., 2025). Current models of self-evaluation focus almost exclusively on the waking mind. Yet, whether sleep, as a critical window for consolidation daytime experience, can strengthen positive self-evaluation remains largely unknown. Addressing this question is crucial not only for understanding how positive self-beliefs can be fostered, but also for explaining why such beliefs may be difficult to strengthen in individuals with elevated depressive symptoms. Here, we investigate how reactivating positive self-evaluative traits during sleep would support self-evaluation and the underlying neurocognitive mechanisms, and to what extent this mechanism may become maladaptive with elevated depressive symptoms.

Indeed, while sleep’s role in memory consolidation is well-established, how it contributes to self-evaluation remains largely unexplored. According to the active systems consolidation account, hippocampus-dependent memory traces of recent experience are repeatedly reactivated during non-rapid eye movement (NREM) slow-wave sleep, with the reactivation gradually consolidating memory traces into stable and neocortex-dependent memory representations (Brodt et al., 2023; Diekelmann & Born, 2010; Lutz et al., 2026). This systems-level consolidation process was coordinated via the hippocampus-neocortex interaction, mediated by the coupling among neocortical slow oscillations (SOs), thalamic sleep spindles and hippocampal ripples rhythms (Staresina et al., 2015, 2023; Ng et al., 2025; Schreiner et al., 2024). Although this framework has been developed primarily in the context of episodic memories, memories about self, particularly self-evaluative episodes, may also require sleep-mediated reactivation and consolidation to stabilize.

Notably, reactivation not only stabilizes memories but also enables adaptive memory transformation for cognitive and emotional functions (Paller et al., 2021; Xia & Hu, 2026). Specifically, sensory cueing during sleep can selectively bias reactivation processes to enhance memory, support value-based decision-making, alleviate clinical symptoms, and promote positive affect, among other benefits (Xia et al., 2024; Z. Yao et al., 2024; Chen et al., 2024; Heijden et al., 2024; Recher et al., 2024; Xia et al., 2023; Pereira et al., 2022; Hu et al., 2020, 2015; Cairney et al., 2014; Rudoy et al., 2009; Rasch et al., 2007). Mechanistically, examining the cue-triggered neural activity has begun to reveal the neurocognitive mechanisms that mediate memory reactivation and cross-regional interaction during consolidation and transformation (Cairney et al., 2018; Duan et al., 2025; Liu et al., 2023; Nicolas et al., 2025; Schreiner et al., 2024). However, prior work has largely focused on neural activation strength or pattern reinstatement per se, rather than whether sleep cueing can organize functionally meaningful neural representations that align with downstream cognition and behavior, i.e., self-evaluation.

Here, we aimed to bridge this gap by unraveling how neural representational structures during sleep would align with post-sleep self-evaluative structures. Specifically, we addressed the following questions. First, whether cueing positive traits during sleep can elicit fine-grained, trait-specific neural representations of the self, rather than global neural activities changes. Second, the extent to which these neural representations map onto post-sleep self-evaluation, i.e., the neural-evaluative representational alignment. Third, given that reduced positive self-evaluation is a core feature of depressive symptomatology, an important open question is whether this neural–evaluation alignment may vary as a function of individual differences in depressive symptoms.

To address these questions, we combined self-evaluation assessments, sleep reactivation and cross-modal representational similarity analyses (RSA). Participants first completed a self-evaluation task to characterize self-evaluation before sleep. During subsequent slow-wave sleep, we re-played half of the positive traits to selectively reactivate positive self-evaluative processing while neutral adjectives served as control cues to isolate the self-evaluative processing. Critically, we employed time-resolved RSA on cue-elicited EEGs to delineate the emergence of trait-specific neural representations of positive self. We next quantified the correspondence between neural representational structure during sleep and post-sleep self-evaluative structures, and examined how such neural–evaluative alignments varied across individuals with varying levels of depressive symptom severity. By constructing individualized representational dissimilarity matrices, we can illuminate whether sleep reactivation organizes fine-grained neural representations in a manner that carries functional significance for post-sleep self-evaluation (Kriegeskorte et al., 2008; Nili et al., 2014; Weber et al., 2024).

## Results

### Sleep reactivation increased categorical positive self-evaluation, with benefits attenuated by depressive symptom severity

We recruited 65 participants spanning a range of depressive symptoms (Beck Depression Inventory BDI-II raw scores: Mean = 13.11, s.d. = 9.71, range = 0 - 37). Participants completed a series of self-referential encoding and self-evaluation tests (Fig. 1a) and underwent targeted memory reactivation (TMR) of positive traits during subsequent slow-wave sleep (Fig. 1c). We treated binary endorsement of positive traits as the primary index of self-evaluation, indicating whether a positive trait was categorically incorporated into the participant’s self-concept (Fig. 1b, upper panel). Continuous descriptiveness ratings were used primarily to define graded self-evaluative structure for representational analyses (Fig. 1b, lower panel). Because only positive traits were cued during sleep, all TMR analyses focused on positive traits. Bayesian multilevel mixed models included TMR condition (cued vs. uncued), Time (post-sleep vs. delayed), standardized BDI-II score (depression severity) modelled continuously, and their interactions, while controlling for pre-sleep item-level responses, age and gender. Conditional estimates at BDI-II z = −1, 0 and +1 are reported only to display the continuous moderation effect rather than separating participants into sub-groups.

**Fig. 1:**
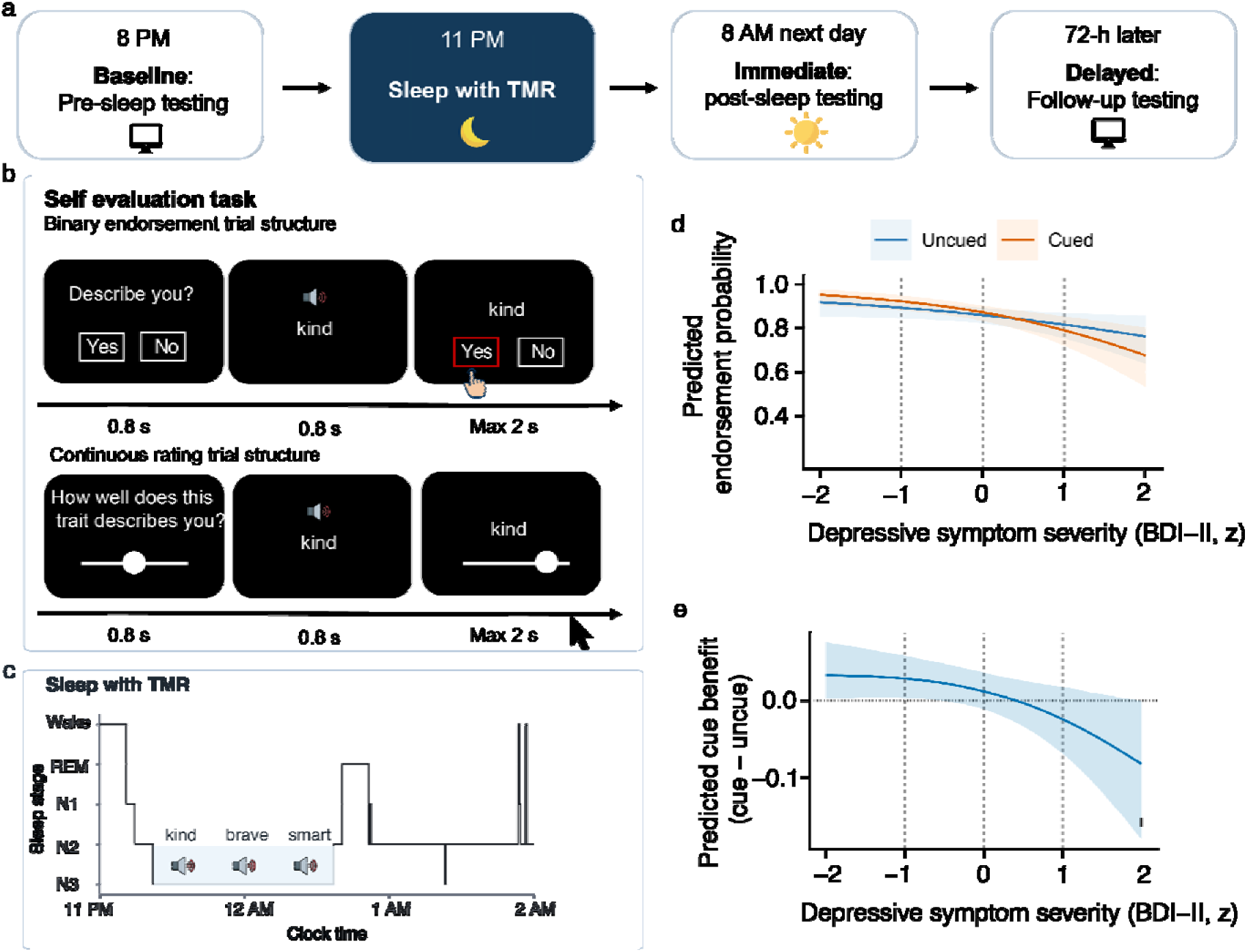
Experimental procedure, task structure and behavioral results. **a,** Experimental procedure. Participants (**N = 65**) completed a pre-sleep self-evaluation test, followed by overnight sleep with targeted memory reactivation (TMR), immediate post-sleep self-evaluative tests the next morning and a 72-h delayed follow-up. During stable slow-wave sleep, half of positive traits was replayed acoustically while EEG was recorded. Depressive symptom severity was measured using the Beck Depression Inventory-II (BDI-II, Beck et al., 1996). **b,** The self-evaluation task. Participants evaluated 40 positive and 40 negative personality traits. For each trait, they made a binary endorsement judgement indicating whether the trait described the self (upper panel), and a continuous descriptiveness rating indicating how well the trait described the self (lower panel). These two measures indexed categorical endorsements and continuous ratings of self-evaluation, respectively. Positive and negative traits were assessed in the self-evaluation task. The primary TMR contrast was restricted to positive traits because only positive traits were replayed during sleep. Additional task measures are reported in the Supplementary Note 2-3. **c,** Illustration of sleep TMR procedure from a representative participant. The hypnogram shows sleep stages from 11 pm to 2 am, during which spoken positive traits from the self-evaluation task were delivered during slow-wave sleep. A separate set of neutral adjectives, familiarized before sleep through valence and familiarity ratings, was replayed as control cues to isolate the self-evaluative processing (see Methods). Cued and uncued positive traits were assigned within participant and were matched based on pre-sleep measures (see Supplementary Note 4). **d,** Model-estimated endorsement probability for cued and uncued positive traits as a function of standardized depressive symptom severity. **e,** Estimated cueing benefit for endorsement of positive traits, calculated as cued minus uncued endorsement probability. Vertical dashed lines mark −1 s.d., the mean and +1 s.d. of BDI-II score; horizontal dotted lines indicate no cueing benefit. Cueing benefits on categorical endorsement were most evident at lower depressive symptom severity.

For binary endorsements of positive traits, we found that the cueing benefits depended on depressive symptom severity, as indicated by a TMR by depression interaction (median = 0.054, 95% CrI [0.005, 0.109], pd = 98.3%, Fig. 1d). Averaging across the post-sleep and delay tests, cueing increased endorsements of positive traits at lower depressive symptom severity (−1 s.d.; cued minus uncued: 2.9 percentage points, 95% CrI [0.2, 5.8], pd = 98.4%), whereas effects were small and uncertain at the mean level of depressive symptoms (1.2 percentage points, 95% CrI [−1.2, 3.6], pd = 84.4%) and showed no clear and even negative effect at higher depressive symptom severity (+1 s.d.; −2.5 percentage points, 95% CrI [−6.9, 1.9], pd = 88.1% in the negative direction). This depression moderation effect was mostly evident during post-sleep immediate test (lower vs. higher depression, median = 0.075, 95% CrI [0.011, 0.147], pd = 98.7%), and became weaker at delay (median = 0.033, 95% CrI [−0.034, 0.108], pd = 83.0%), although the direct TMR × depression × Time interaction contrast was negative and uncertain (Delay minus Post-sleep: median = −0.042, 95% CrI [−0.133, 0.049], pd = 82.1%). Together, these results indicate that sleep reactivation elevated categorical endorsement of positive self-traits among individuals with lower depressive symptom severity.

In addition to binary endorsement, participants also provided continuous descriptiveness ratings of positive traits. These continuous ratings were used to characterize graded item-level structure of self-evaluation for the neural-evaluative representational analyses. Full behavioural analyses of descriptiveness ratings are reported in Supplementary Note 1.

### Cueing elicited robust EEG spectral power changes but showed limited links to depressive symptom severity and behavioural updating

We first confirmed that replaying positive traits during slow-wave sleep elicited typical EEG signatures of cue processing. Self-evaluative neural responses were quantified as the difference in time-frequency-resolved EEG power between trait cues and control cues (Trait minus Control). Cluster-based permutation testing identified two significant cue-elicited EEG power clusters (cluster-level *p* < 0.05; Fig. 2a): an early 2.00–11.50 Hz cluster from 0.260 to 0.940 s, and a later 9.50–21.00 Hz cluster from 0.700 to 1.750 s.

**Fig. 2:**
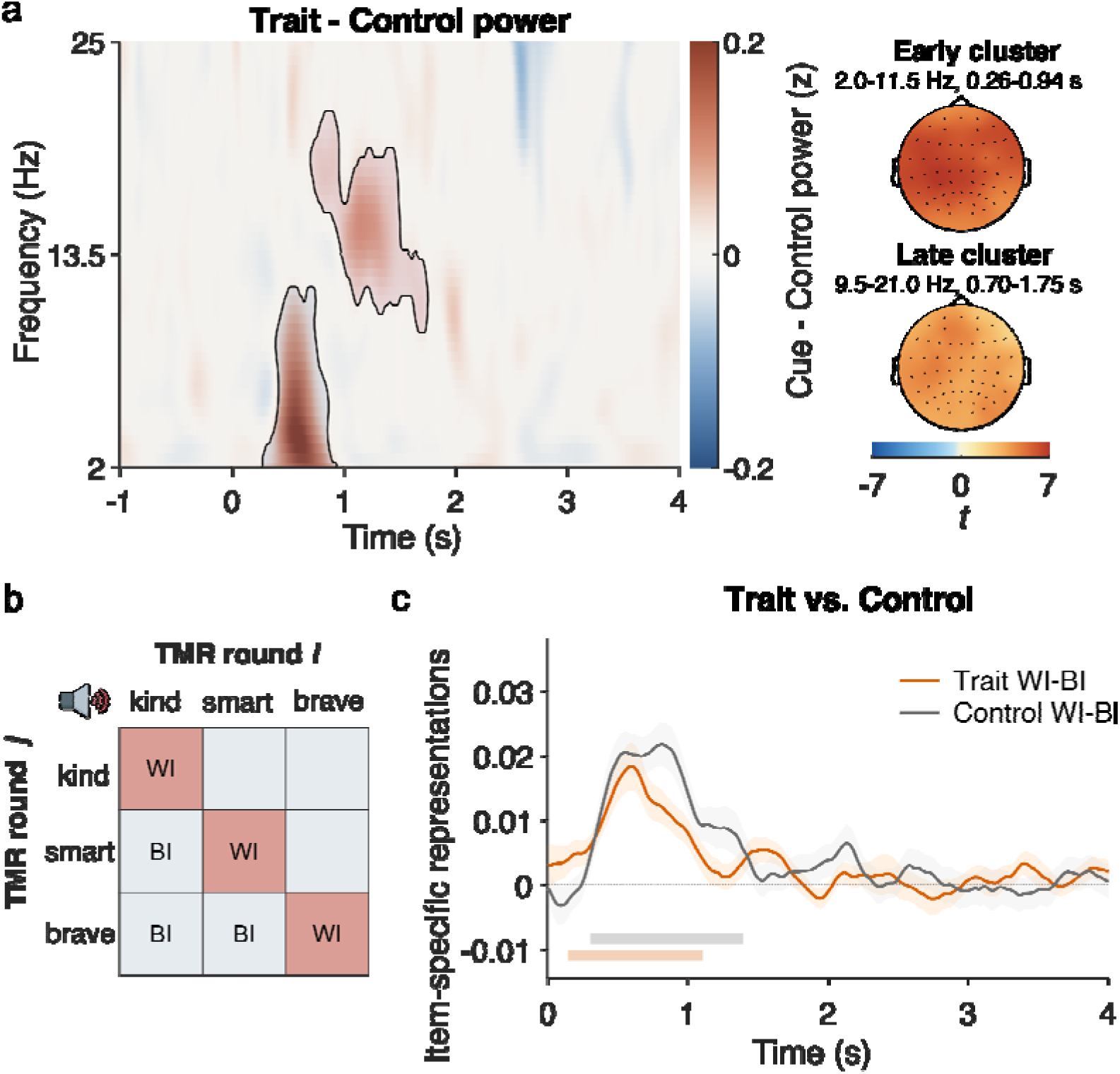
Cueing during sleep elicits oscillatory power changes and item-specific neural representations. **a,** Time–frequency representations of the positive trait cues minus control cue EEG power contrast during NREM sleep. Black contours outline significant clusters identified by cluster-based permutation testing. Scalp maps show the corresponding topographies of the early cluster (2.0–11.5 Hz, 0.26–0.94 s) and late cluster (9.5–21.0 Hz, 0.70–1.75 s), displayed as *t* statistics. Scalp maps show the corresponding topographic t statistics averaged within the corresponding cluster windows: early cluster, 2.0–11.5 Hz, 0.26–0.94 s; late cluster, 9.5–21.0 Hz, 0.70–1.75 s. **b,** Schematic of the raw EEG representational similarity analysis (RSA). Item-specific neural representations were quantified by comparing similarity for repeated presentations of the same trait across TMR rounds (within-item, WI) with similarity for different traits across rounds (between-item, BI). **c,** Time-resolved raw EEG RSA across all participants, shown separately for cue and control trials. The plotted index reflects item-specific neural representation strength, quantified as the difference between within-item and between-item similarity (WI − BI). Higher values indicate stronger item-specific representations. Solid lines denote group means, and shaded ribbons denote ± s.e.m. Horizontal bars indicate significant time clusters from one-sample cluster-based permutation tests performed separately for cue and control trials.

We then asked whether these cue-elicited power responses were related to depressive symptom severity or subsequent behavioural updating. Mean Trait − Control power extracted from the significant group-level clusters showed no reliable association with BDI-II score after controlling for age and gender, and item-level power did not provide reliable evidence for predicting post-sleep endorsement or for power × BDI-II moderation. Full participant-level permutation regressions and Bayesian neural-evaluative models are reported in Supplementary Note 5.

Together, these results demonstrate that re-playing positive traits elicited robust cue-elicited EEG responses during sleep. While EEG power changes indicated the level of cue processing or strength of reactivation, they do not reveal the trait-specific representational structures. Therefore, we next examined whether cue-elicited EEG would contain trait-specific representational structure using representational similarity analyses (RSA).

### Cueing elicited early item-specific EEG patterns during sleep

We next tested whether cue presentations elicited item-specific EEG patterns during sleep. Sixty-two participants were included in the item-level multivariate EEG RSA (three additional participants were excluded because they contributed fewer than three clean TMR blocks due to excessive artifacts). Item-level similarity was quantified as the difference between within-item and between-item multichannel EEG similarity using only cross-block trial pairs, thereby minimizing temporal proximity confounds (Fig. 2b). Across participants, both cued and control items showed greater within-item than between-item similarity during early post-cue intervals (cue: 0.14–1.10 s, cluster-corrected *p* < 0.001; control: 0.30–1.39 s, cluster-corrected *p* < 0.001; Fig. 2c), indicating item-specific EEG patterns during sleep. However, positive trait cues and control adjectives did not significantly differ from each other in item-level similarity across the epoch (minimum cluster-level *p* = 0.17). Participant-level item-specific EEG similarity was not reliably associated with BDI-II score after controlling for age and gender, and item-level similarity did not provide reliable evidence for predicting post-sleep endorsement or for similarity × BDI-II moderation. Thus, early item-specific EEG patterns may reflect general stimulus discrimination during sleep. Full results are reported in Supplementary Note 5.

### Neural-evaluative alignments vary with depressive symptoms

We next asked our key question: to what extent item-specific neural representations during sleep would align with the post-sleep self-evaluative representations, and whether this neural– evaluative correspondence may vary with depressive symptoms. Because continuous descriptiveness ratings provide a graded, item-level measure of self-evaluation, we constructed self-evaluative representational dissimilarity matrix (RDM) based on the pairwise differences in descriptiveness ratings averaged across the post-sleep immediate and delayed tests (Fig. 3a). This averaged RDM was used to obtain a stable estimate of post-sleep self-evaluative representational structure; sensitivity analyses using immediate or delayed ratings yielded similar neural– evaluation alignment patterns (Supplementary Fig. 1).

**Fig. 3:**
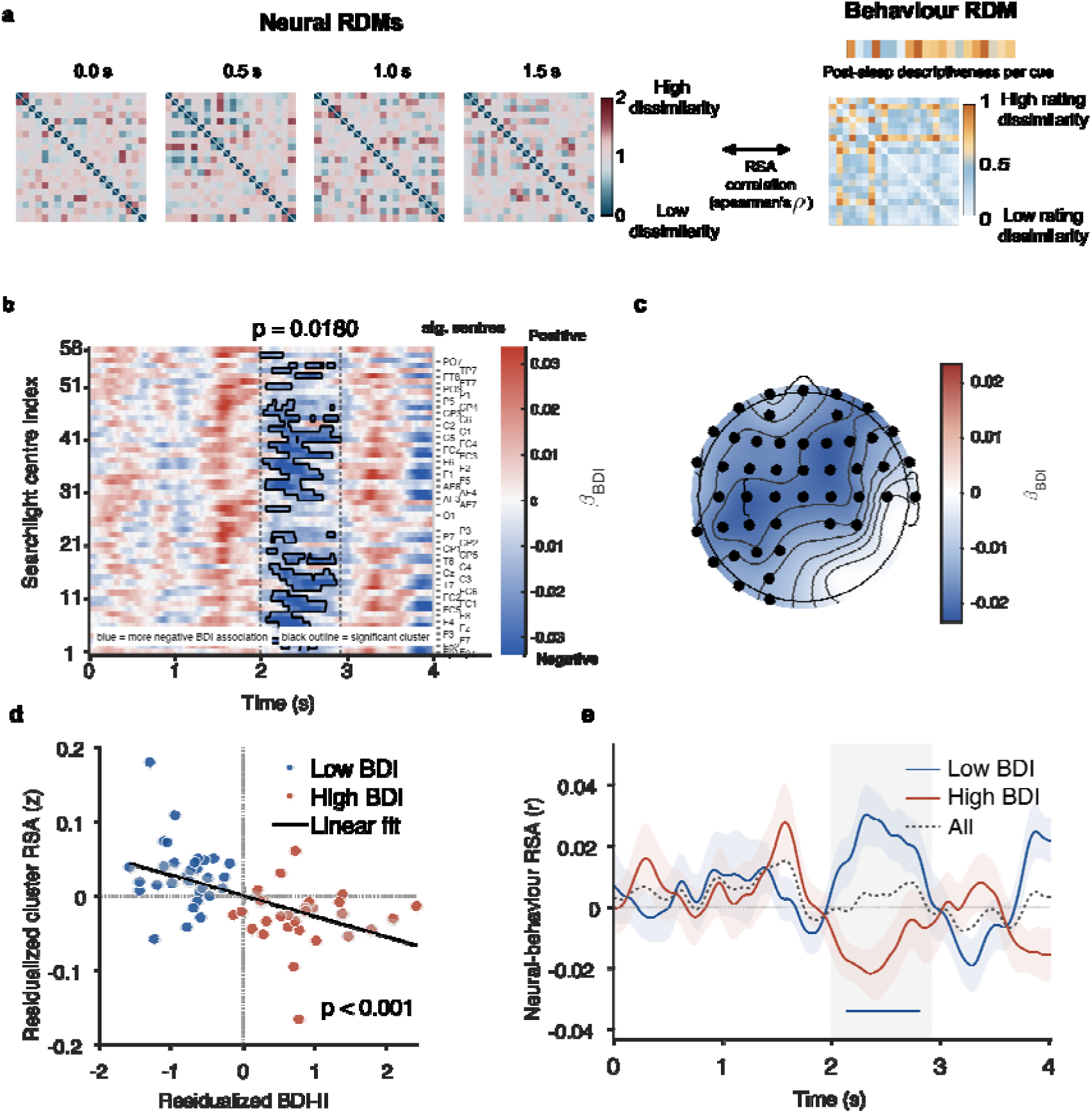
Depressive symptom severity was associated with reduced late neural–behaviour representational alignment during sleep. **a,** Searchlight representational similarity analysis. Neural RDMs were computed from cue-elicited 11–16 Hz EEG power in 500-ms sliding windows advanced in 10-ms steps across the 0– 4 s post-cue interval and correlated with a self-evaluative RDM derived from post-sleep descriptiveness ratings averaged across the post-sleep and delayed assessments. **b,** Sensor × time map of the BDI-II regression effect on neural–evaluative RSA alignment. Colours indicate the BDI-II regression coefficient after adjustment for age and gender; blue indicates a stronger negative association between depression and RSA alignment. The black contour denotes the cluster surviving cluster-level permutation correction. **c,** Topography of the significant negative BDI-II searchlight cluster. Colours indicate the depression regression coefficient after adjustment for age and gender; highlighted sensors indicate cluster searchlight centres. **d,** Exact-cluster neural-evaluative alignment plotted against residualized BDI-II score. Colours denote predefined BDI-II sub-groups used for visualization only; while statistical inference was based on the continuous BDI-II depression model. **e,** Time-resolved neural-evaluative alignment by predefined depressive symptom sub-groups. Horizontal bars indicate cluster-corrected positive-alignment windows. Complementary endorsement-based RDMs did not yield cluster-corrected BDI-II depression moderation effects.

Searchlight-based, pairwise neural RDMs were constructed from cue-elicited sigma 11–16 Hz power during slow-wave sleep. We focused on the spindle-related 11–16 Hz sigma power because it is implicated in memory reactivation and particularly item-specific patterns (Antony et al., 2018; Cairney et al., 2018; Liu et al., 2023; Schreiner et al., 2021, 2024). Neural RDMs were computed with a sensor-level spatiotemporal EEG sigma power searchlight to estimate region-specific representational structures. Each searchlight comprised a centre EEG sensor and its four nearest neighbouring sensors, with this procedure being repeated for each of the 58 sensors across successive time windows. Local sigma-power patterns were extracted from 0–4 s after cue onset using 500-ms windows advanced in 10-ms steps. Within each searchlight and time window, neural dissimilarity between trait cues was defined as 1− *r*, where r was the Fisher-z-averaged, back-transformed Spearman correlation across cross-block trial-level sigma-power patterns.

Neural–evaluation alignment was quantified as the Fisher-z-transformed Spearman correlation between vectorized neural and behavioural RDMs. Because our primary hypothesis concerned depression moderation of this alignment, we regressed alignment values at each sensor–time searchlight on standardized BDI-II score, adjusting for age and gender and applying cluster-level permutation correction across sensors and time.

This alignment analysis identified a significant negative association between standardized BDI-II score and neural–evaluative representational alignment. The cluster extended from 1.99 to 2.92 s after cue onset and comprised 47 searchlight centre sensors after adjustment for age and gender (cluster-level permutation *p* = 0.008; Fig. 3b, c). To visualize this negative association, we extracted each participant’s mean RSA value from the significant sensor–time cluster and correlated this value with the continuous BDI-II score (β = −0.0276, *t* = −4.75, *p* < 0.001, covariate-adjusted, Fig. 3d). This alignment-depression association was further highlighted in the lower- and higher-symptom sub-groups, using conventional threshold (BDI-II ≤ 13 versus ≥ 14; see Methods). As expected, this descriptive grouping showed stronger representational alignment in the lower-than in the higher-depression group (BDI ≤ 13: n = 31; BDI ≥ 14: n = 31; Welch t (59.15) = 5.39, *p* < 0.001; Fig. 3e). Moreover, in the lower-depression group, RSA within the exact cluster was significantly greater than zero (mean Fisher-z RSA = 0.0312, t = 4.00, *p* < 0.001), whereas the full sample did not show a significant positive alignment effect (*p* = 0.325). Supplementary analyses applying the same searchlight RSA procedure to theta and beta bands did not reveal depression-related alignments (Supplementary Note 6 and Supplementary Fig. 2), suggesting that the effects were specific to sigma activity.

Because the effect occurred in the spindle-related 11–16 Hz sigma range, we further conducted a descriptive midline-overlap follow-up analysis to assess whether the depression-related neural– evaluation alignment was preserved at frontocentral sensors commonly used to characterize NREM sigma and spindle activity. This follow-up ROI was restricted to searchlights centred on Fz, FCz and Cz that overlapped the primary 47-sensor searchlight cluster. In this 3-sensor midline ROI, the negative alignment-depression association was reproduced (β = −0.0300, *t* = −3.54, *p* < 0.001), and a time-resolved follow-up showed a cluster-corrected negative alignment-depression association from 2.20 to 2.57 s after cue onset (*p* = 0.037). Additional descriptive checks using lower- and higher-symptom sub-groups and positive-alignment windows are reported in the Supplementary Information (Supplementary Note 7). Together, these results indicate that trait-specific neural representational structure during sleep aligned with the post-sleep self-evaluative representational structure, with this alignment being most evident among individuals with lower depressive symptom severity.

### Depression moderated how spindle predicted post-sleep positive self-evaluation

Given that the sigma activity supports the neural-evaluative representational alignment, we next performed a targeted spindle event analysis to investigate how spindle may predict post-sleep positive self-evaluation and how depression may moderate this prediction. We used YASA to derive baseline-corrected cue-elicited spindle probability averaged across midline RSA sensors (Fz, FCz and Cz) across the 0–4 s post-cue interval (Fig. 4a and 4b, see Methods). This analysis revealed significant positive spindle probability clusters for both trait and control cues. Replaying trait cues significantly increased spindles from 1.08 to 2.06 s after cue onset, whereas control cues significantly increased spindles from 0.94 to 2.15 s (all *p*s < 0.001, Fig. 4c). Thus, both cues elicited robust spindle-probability responses in slow-wave sleep.

**Fig. 4:**
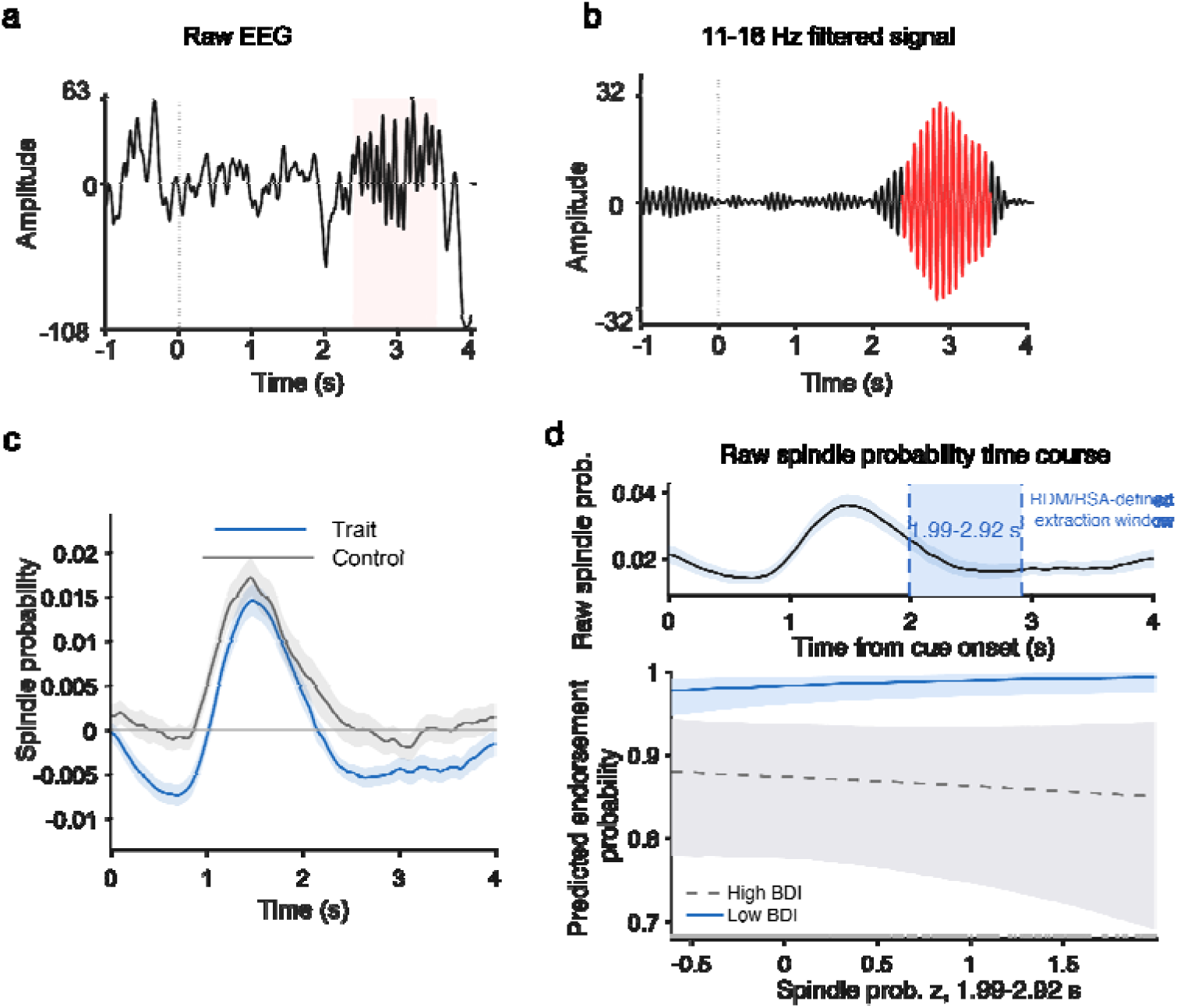
Cue-elicited spindle probability and its relationship to self-evaluations. **a**, Example single-trial N3 EEG trace averaged across Fz, FCz and Cz from −1 to 4 s relative to cue onset. The shaded interval marks the YASA-detected spindle event shown in **b**. **b**, The same EEG segment after 11–16 Hz band-pass filtering. The red segment indicates the detected spindle event. **c**, Baseline-corrected cue-elicited spindle probability averaged across Fz, FCz and Cz for positive trait cues and neutral control cues. Lines show the group mean for trait cues and control cues; shaded areas indicate ±s.e.m. Spindle probability was baseline-corrected relative to the pre-cue interval. Color matched horizontal bars indicate cluster-corrected post-cue intervals in which spindle probability significantly increased relative to the pre-cue baseline: trait cues, 1.084–2.036 s, cluster-level *p* = 0.0003999; control cues, 0.956–2.116 s, cluster-level *p* = 0.0002. **d**, Top, cue-elicited spindle probability for positive traits; blue shading marks the independently defined sigma neural-evaluative alignment window (1.99–2.92 s post cue). Bottom, model-predicted post-sleep endorsement probability as a function of z-scored spindle probability extracted from this window, shown at lower and higher BDI-II depression levels. Shaded bands indicate 95% credible intervals; rug marks show the observed spindle probability distribution.

To test whether depressive symptoms were associated with the magnitude of the cue-elicited spindle probabilities, we extracted mean baseline-corrected spindle probability from the significant group-level spindle window and related it to BDI-II score using participant-level covariate-adjusted permutation regression. Results revealed that the BDI-II score did not reliably predict this baseline-corrected spindle probabilities after controlling for age and gender.

For behavioural prediction, we used baseline-corrected post-cue spindle probability in the RSA-aligned fixed spindle window (i.e., 1.99−2.92 s), and included pre-cue baseline spindle probability as an additional covariate. In this Bayesian mixed-effects logistic regression model, the spindle probability × depression interaction was negative, posterior mean = −0.292, 95% CrI [−0.583, −0.015], pd = 98.0% (Fig. 4d). Consistent with this interaction, higher spindle probability positively predicted endorsements of positive traits at lower depressive symptom severity, posterior mean = 0.481 on the log-odds scale, 95% CrI [0.003, 0.999], pd = 97.6%, whereas this association was small and uncertain at higher depressive symptom severity, posterior mean = −0.103, 95% CrI [−0.425, 0.218], pd = 73.1% in the negative direction.

Taken together, sleep cueing of positive traits elicited robust univariate EEG responses and significant item-specific EEG patterns. Most importantly, sigma power contained trait-specific neural representations during sleep aligned with post-sleep self-evaluative representational structures, with the strength of alignment being strongest among individuals with lower depressive symptoms and attenuated as elevated depressive symptom severity.

## Discussion

The present study identifies a sleep-based representational mechanism through which positive self-evaluation may be strengthened and shows that this mechanism is weakened with increasing depressive symptom severity. Reactivating positive personality traits during NREM sleep promoted categorical endorsement of positive traits as self-descriptive, with the benefit most evident among participants with lower depressive symptoms. At the neural level, positive-trait cues elicited robust cue-elicited EEG responses and early item-specific neural patterns during sleep. Critically, however, the neural signature most closely linked to self-evaluation was not the overall magnitude of cue-elicited activity nor early item-specificity, but a later sigma-band representational structure that aligned with the graded structure of post-sleep self-evaluation.

Across a large sample of participants with varying levels of depressive symptoms, we further demonstrated that this neural-evaluative alignment was attenuated as depressive symptoms increased. Together, these findings suggest that sleep reactivation may strengthen positive self-evaluation when reactivated neural patterns preserve behaviorally meaningful self-evaluative structure, whereas elevated depressive symptoms are associated with reduced neural alignment of reactivated positive self-representations.

Cueing positive traits during NREM sleep enhanced positive self-evaluation, as indicated by increased endorsement of cued positive traits, consistent with prior research documenting the behavioral benefits of targeted memory reactivation (Cairney et al., 2014; Groch et al., 2017; Heijden et al., 2024; Hu et al., 2020; Whitmore et al., 2022; Xia et al., 2024; Z. Yao et al., 2024). Electrophysiologically, replaying self-evaluative positive traits reliably elicited EEG spectral responses during sleep, indicating self-evaluative processing. However, the magnitude of these univariate neural responses showed limited evidence of behavioural relevance and did not account for depressive symptom-related differences in self-evaluation changes.

Indeed, our primary result went beyond univariate modulation of neural activities: instead, we found that the searchlight-based, trait-specific neural representational structure during sleep reactivation was strongly aligned with the post-sleep self-evaluative representational structure, with the strength of the alignment inversely tracking depressive symptom severity. Temporally, this depression-related neural–evaluation alignment was most evident in a late window from 1.99 to 2.92 s post cue onset. These findings were further complemented by a midline-restricted analysis that identified an overlapping temporal cluster from 2.2 to 2.57 s. The temporal dissociation between early item-specific neural representations and later sigma-mediated neural– evaluative alignment suggests a functionally dissociable, sequential memory processing (see also (Antony et al., 2019; Duan et al., 2025; Lewis & Bendor, 2019; Liu et al., 2023). Early after cue onset, both trait and control cues elicited item-specific neural representations, suggesting an initial sensory processing and discrimination between individual auditory cues that was not, by itself, sufficient to account for later post-sleep changes in self-evaluation. By contrast, the neural signature that tracked post-sleep self-evaluative representations emerged later, within a sigma-band window centered on approximately 2.0–3.0 s after cue onset. This later sigma-based representational structure may thus reflect reactivation of self-evaluative processing that was further integrated into the self-evaluative schema, improving post-sleep positive self-evaluations. This interpretation is supported by converging evidence linking post-cue spindle-related neural activity in late windows to successful memory reactivation (Antony et al., 2018; Cairney et al., 2018; Duan et al., 2025; Liu et al., 2023; Schreiner et al., 2024).

Sleep spindles are canonical physiological markers of NREM memory reactivation and provided a complementary, event-level perspective on the sigma-band alignment findings. Prior research establishes that post-cue spindle activity and slow-oscillation–spindle complexes are conducive to successful reactivation, item-specific memory representation and predicted memory retention (Antony et al., 2018; Cairney et al., 2018; Liu et al., 2023; Schreiner et al., 2021, 2024). Whereas RSA captured multivariate structure in cue-elicited sigma-band activity (11–16 Hz), the spindle analysis tested whether this same temporal window contained discrete sleep markers with behavioral significance. We found that spindle probability within this RSA-defined window predicted categorical positive self-endorsement, again in a depression-dependent manner with the strongest association observed in individuals with lower depressive symptoms. Thus, these results link multivariate representational structure to a canonical sleep oscillation implicating memory reactivation, providing mechanistic insights that the fine-grained neural–evaluation alignment may be grounded by spindle-mediated reactivation dynamics.

Beyond demonstrating a neural-evaluative alignment during sleep, our results advanced the understanding of depression-related cognition and how intrinsic motivational states (e.g., depression) can interact with sleep reactivation. Depression is marked by low self-worth and reduced positive self-referential processing: individuals with higher depressive symptoms tend to endorse fewer positive traits and show broader biased in negative self-evaluative processing (Collins & Winer, 2024; Dainer-Best et al., 2017; Disner et al., 2011; Dozois & Beck, 2008). Our findings provide a new mechanistic account to explain this maladaptive self-evaluation patterns. While participants with higher depressive symptoms still showed early item-specific neural representations implicating auditory processing of individual cues, the late sigma-band neural structure was less coupled with graded self-evaluative structures (Collins & Winer, 2024; Pizzagalli, 2014). Such weakened coupling may help explain why sleep reactivation of positive traits was insufficient to enhance categorical positive self-endorsement at higher depressive-symptom severity. This perspective also joins recent evidence suggesting that emotional, motivational and reward-related factors can bias both endogenous and exogenous sleep-dependent memory reactivation and consolidation (Davidson et al., 2021; Igloi et al., 2015; Wilhelm et al., 2011), highlighting that intrinsic motivational and affective states represent key modulators of the representational structures and outcomes of sleep memory reactivation (Hu et al., 2020; Oudiette & Paller, 2013).

Limitations and future directions should be noted. First, the present sample varied in depressive symptom severity but did not consist of clinically diagnosed participants with major depressive disorder. Therefore, the findings should be interpreted as evidence about dimensional depressive symptoms rather than clinical depression. Future studies in diagnosed clinical populations should test whether sigma-band neural-evaluative alignment predicts clinically relevant self-evaluative changes, and whether enhancing spindle-related reactivation can strength positive self-endorsement (Jourde et al., 2025; Kiang et al., 2017; Mutreja et al., 2026; Shin et al., 2025; J. Yao et al., 2023). Second, while the multivariate RSA revealed fine-grained item-specific neural representations, the underlying precise neural circuits remains unclear. Future work shall employ intracranial EEG or multimodal imaging to test how hippocampal-neocortical interactions support the reactivation and integration of positive self-evaluative information (Abdellahi et al., 2026; Duan et al., 2025; Schreiner et al., 2024).

In sum, targeted reactivation of positive traits during sleep selectively strengthens positive self-evaluation, with this benefit attenuated as depressive symptoms increase. Mechanistically, reactivation elicited trait-specific neural representations during sleep that mapped onto post-sleep self-evaluation, which spindle-related sigma activity supporting this personalized, fine-grained alignment. Critically, this representational alignment was disrupted in individuals with elevated depressive symptoms, potentially contributing to the well-documented rigidity and diminished positivity of self-evaluation. Together, these findings identify the sleep-based, personalized neural-evaluative alignment as a candidate mechanism through which reactivation shapes self-evaluation, and offers a new explanation for why positive self-views are less readily strengthened in individuals with elevated depressive symptoms. More broadly, our findings establish a new approach for understanding, and potentially modifying the neural architecture of the self in health and mental disorder.

## Methods

### Participants

The final sample included 65 adults with usable behavioural and EEG data for analyses (8 male; mean ± SD age = 22.7 ± 2.5 years). Participants were recruited to span a broad range of depressive symptom severity using Beck Depression Inventory–II (BDI-II) in order to test whether depressive symptoms moderate the behavioral and neural effects of overnight TMR. All primary inferential analyses treated BDI-II score as a continuous variable to capture the full variances of the sample. For descriptive purposes only, we used conventional BDI-II cutoff to divided participants into a lower-symptom sub-group (BDI-II 0–13; n = 33; mean ± s.d. = 5.0 ± 4.3; range, 0–13) and a higher-symptom sub-group (BDI-II ≥ 14; n = 32; mean ± s.d. = 21.4 ± 5.7; range, 14–37). To the best of our knowledge, this sample size was much higher than recent sleep and reactivation studies, which typically recruited 20-40 participants (Chen et al., 2024; Schechtman et al., 2023; Xia et al., 2023; Z. Yao et al., 2024). To maximize the likelihood of successful cue delivery during stable slow-wave N3 sleep, eligibility required a habitual 7–9 h sleep window between 00:00 and 09:00 without severe insomnia symptoms (Insomnia Severity Index, ISI ≤ 14). Exclusion criteria included a history of psychiatric, neurological, or sleep disorders, colour vision deficiency, current medication use, and alcohol or caffeine consumption on the day prior to or the day of testing. An additional thirteen participants were excluded prior to analysis: insufficient slow-wave sleep (n = 9); heard word during TMR (n = 3, see ‘Targeted memory reactivation’ below); or not native Mandarin speaker (n =1). All participants provided written informed consent, received HKD 500 (≈ USD 64) upon completion, and the protocol was approved by the Human Research Ethics Committee, The University of Hong Kong.

### Stimuli

The stimulus set comprised 80 personality-trait adjectives (40 positive, 40 negative) and 10 neutral adjectives used as control cues (e.g., positive: “competent”; negative: “selfish”; neutral: “slender”). All items were presented visually and aurally during behavioural testing. For sleep cueing, auditory recordings were prepared for the positive traits selected as TMR cues and for the neutral control adjectives. The full stimulus list is provided in the Supplementary Table 1.

### Procedure and Tasks

Participants completed two laboratory visits separated by approximately 72 h. During visit 1 (start time 19:30–20:30), participants provided informed consent, completed questionnaires, and performed the Psychomotor Vigilance Task (PVT) to index alertness. They then completed pre-sleep behavioural testing, followed by overnight EEG recording with TMR delivered during stable N3 sleep. Upon awakening, participants completed immediate post-sleep testing (post-sleep). During visit 2 (∼72 h later), participants completed delayed testing (Delay) (Fig. 1). All tasks were implemented in PsychoPy (v2020.1.3).

### Questionnaires

During visit 1, participants completed a series of questionnaires assessing self-esteem, personality traits and emotion regulation styles, etc (for a full list of questionnaires and descriptive statistics, see Supplementary Table 2). These questionnaires served as a cover story to support engagement with the subsequent self-evaluative tasks and were not analysed.

### Psychomotor vigilance task PVT

Participants completed a 5-min PVT before each behavioural session (pre-sleep, post-sleep and delay) to assess vigilance. On each trial, a central fixation cross was presented for a jittered interval of 2–10 s, followed by a counter starting from 0; participants responded as quickly as possible when the counter appeared, and reaction time (RT) was displayed as feedback. Vigilance was indexed as the median RT across valid trials for each participant and session. A one-way repeated-measures ANOVA on median RTs revealed no significant effect of Session, F(2, 88) = 2.09, p = 0.13, indicating comparable vigilance across sessions.

### Self-evaluation tasks and primary outcomes

At pre-sleep baseline self-evaluation test (adapted from the self-referential encoding task, SRET), participants were presented with 80 personality trait adjectives (40 positive, 40 negative) and made speeded yes/no decisions indicating whether the trait would describe themselves. On each trial, a trait word was presented for 800 ms, followed by a 2 s response window. Responses were made using the left and right arrow keys, with response mapping counterbalanced across blocks. The inter-trial interval varied randomly between 1.2 and 1.4 s.

Immediately following the binary endorsement task, participants completed a continuous rating block involving the same 80 trait adjectives, presented again in the same order. For each word, participants rated its self-descriptiveness and importance to self-identity. Each trait was displayed for 800 ms, and participants had up to 8 s to provide their ratings on a continuous rating bar ranging from 0 to 100. This rating block increased the salience of the traits prior to sleep, and yielded item-level self-descriptiveness ratings that were subsequently used to select individualized cue traits for TMR.

After completing the self-evaluative ratings, participants rated 80 personality traits and 10 neutral adjectives on familiarity and emotional valence on continuous scales ranging from 0 to 100. Each word was presented for 800 ms, and participants were allowed up to 8 s to respond. This task was included to further strengthen the semantic and affective representations of all stimuli prior to sleep. Baseline testing concluded with a free-recall task, in which participants were given 5 min to type as many traits as they could remember from the preceding tasks.

The primary positive self-evaluation index was trial-level binary endorsements of positive traits. Continuous self-descriptiveness ratings were used to construct a personalized, graded item-level self-evaluative structure for subsequent neural-evaluative representational similarity analyses. Importance rating and free-recall performance were used to characterise encoding strength and memory for the traits. Secondary behavioural analyses of self-descriptiveness ratings, importance ratings and recall are reported in Supplementary Note 1-3 and Fig. 2-4.

### Traits selection for sleep reactivation

The trait assignment was individualized using baseline self-descriptiveness ratings from the continuous rating block. For each participant, the 40 positive traits were rank-ordered by self-descriptiveness. Twenty traits were then selected as TMR cues using a predefined counterbalanced sampling scheme that spanned the full rating distribution while avoiding adjacent items, thereby minimizing systematic baseline differences between cued and uncued positive traits. The remaining 20 positive traits served as uncued items. In addition, 10 neutral adjectives from the previous familiarity/valence rating task served as control cues during sleep.

### Targeted Memory Reactivation (TMR) during NREM Sleep

Following completion of the pre-sleep behavioural tasks, participants were prepared for overnight polysomnographic recording and slept in a quiet, darkened laboratory bedroom. Continuous white noise (approximately 38 dB) was delivered via a loudspeaker positioned above the bed to minimize stimulus-induced arousals and mask environmental sounds. Throughout the night, trained experimenters monitored the EEG and auxiliary channels (F3/4, FC3/4, C3/4, EOG, EMG) in real time. TMR was initiated only after stable slow-wave N3 sleep, which was visually identified for at least 2 min.

During TMR, auditory presentations of positive trait adjectives (∼40 dB) were delivered through the same speaker, superimposed on the white-noise background. Cueing was restricted to stable slow-wave and N2 sleep and was immediately suspended if signs of arousal or transitions to N1, wakefulness, or REM sleep were detected. Each TMR block comprised 20 positive traits and 10 neutral control words. Each word was presented for ∼1 s, with a randomized inter-stimulus interval of 5–6 s, and successive blocks were separated by a 30 s rest period. Stimulation continued until 02:00 or until 30 cueing blocks had been completed, whichever occurred first.

An a priori inclusion criterion required participants to complete at least four TMR blocks, corresponding to a minimum of 80 positive trait presentations, to ensure sufficient data for EEG analyses. Participants were allowed to sleep for 8 h. If participants were in REM sleep or N3 at the scheduled wake time, awakening was delayed until a transition to N1 or N2 was observed. Upon awakening, participants refreshed themselves, had breakfast, spent approximately 5 min outdoors to reduce sleep inertia, and subsequently reported their subjective sleep quality.

### Post-sleep and Delayed Tests

At both post-sleep and delay tests, free call was assessed before any re-exposure to traits. Participants completed free recall, the binary endorsement self-evaluative task and then the continuous rating task using the same procedures as baseline. The familiarity/valence rating task was not repeated. Participants were not informed in advance about the content or order of the tasks. Participants were debriefed and compensated after completing the study.

### EEG Acquisition and Preprocessing

EEG data were recorded with a 64-channel cap arranged according to the international 10–10 system (eego, ANT Neuro, Enschede, Netherlands) and sampled at 500 Hz. AFz served as the ground electrode and CPz as the online reference, with impedances maintained below 20 kΩ. Bipolar electrooculogram (EOG) electrodes were placed inferior to the left and right eyes, and electromyogram (EMG) electrodes were attached to the chin/jawline to monitor muscle tone. EEG Preprocessing was performed in MATLAB (MathWorks) using custom scripts and the EEGLAB toolbox (Delorme & Makeig, 2004). Continuous overnight EEG recordings were downsampled to 250 Hz, notch-filtered at 50 Hz, and re-referenced to the averaged mastoids. Data were then band-pass filtered 0.5–40 Hz. EOG/EMG were retained for sleep staging but were excluded from the following analyses.

### Sleep Staging

Sleep stages (N1, N2, N3 and REM) were scored from EEG, EOG and EMG using the Yet Another Spindle Algorithm (YASA) with automated staging followed by visual inspection and manual correction where necessary (Vallat & Walker, 2021). Staging was based primarily on electrode C4; when C4 was marked as a bad channel, C3 or Cz was substituted. In accordance with YASA recommendations, EEG data were re-referenced to Fpz for sleep staging. Summary sleep parameters, including total sleep time and stage durations were reported in Supplementary Table 3.

### EEG time-frequency analysis

To characterize neural responses elicited by playing traits during sleep, we analysed cue-elicited EEG activity elicited by positive self-referential trait cues relative to acoustically presented neutral control words. EEG data were epoched from −1.5 to 5.5 s relative to cue onset, allowing analysis of cue-related activity from −1 to 4 s while avoiding edge artifacts. Artifact-contaminated epochs were excluded following visual inspection. Time–frequency decomposition was performed using variable-cycle Morlet wavelets (3–10 cycles) spanning 2–25 Hz in 0.5-Hz steps. Power estimates were baseline-normalised within each experimental block using a pre-cue interval (−1000 to −100 ms), computed separately for each channel and frequency, yielding z-scored power values. Based on prior work on sleep-dependent memory reactivation, cue-related power was quantified as the within-subject difference between positive-trait cues and neutral control cues (Cue − Control). The primary inferential analyses treated BDI-II score continuously; grouped plots were used for visualization only. Time–frequency decomposition and subsequent statistical analyses were implemented using the FieldTrip toolbox (Oostenveld et al., 2011).

### Representational similarity analysis

We conducted two complementary representational similarity analyses (RSA) to address our research questions: whether cueing elicits item-specific neural representation, and whether neural representation would align with post-sleep self-evaluations. The first RSA used raw cue-elicited EEG to quantify item-specific neural representation. The second RSA used cue-elicited sigma-band power to quantify time-resolved correspondence between neural representational structure during sleep and post-sleep self-evaluative rating structure.

### Item-specific neural representations

We first tested whether playing individual traits would elicit item-specific patterns in the cue-elicited EEG signal. Analyses were restricted to the 0–4 s interval following cue onset. EEG epochs were resampled to 100 Hz, and trial-wise response patterns were analysed across all scalp channels. Consecutive 500-ms windows were advanced in 10-ms steps across the analysis interval. Within each window, the neural feature vector for each trial was defined by the concatenated channel × time-sample values.

Similarity between trials was quantified using Spearman correlation. To reduce temporal autocorrelation and shared local-state effects, only trial pairs drawn from different TMR blocks were included. At each time window, within-item similarity was defined as the mean similarity between trials elicited by the same cue across blocks, whereas between-item similarity was defined as the mean similarity between trials elicited by different cues. Item-specific neural pattern strength was quantified as the within-minus-between similarity difference (WI − BI), with similarity estimates Fisher z-transformed before averaging. WI − BI time courses were computed separately for cued items and control items; direct cue-versus-control contrasts were assessed in parallel.

### Searchlight neural-evaluative representational alignment

Neural-evaluative representational similarity analysis tested whether the structure of cue-elicited, item-specific neural patterns during sleep aligned with the structure of post-sleep self-evaluation, and whether this neural-evaluative representational alignment varied with depressive symptom severity across individuals. For the primary analysis, neural RDMs were constructed from cue-elicited 11–16 Hz sigma power during slow-wave sleep. This frequency range was selected because it overlaps canonical sleep spindle frequencies and has been implicated in NREM memory reactivation. The analysis quantified continuous sigma-band power rather than discrete spindle events, allowing us to test whether trait-specific representational structure was expressed in cue-elicited sigma-band EEG activity during sleep. Sigma power was estimated with complex Morlet wavelets at 1-Hz resolution, with cycles increasing linearly from 5 to 8 across 11–16 Hz. Power was sampled every 10 ms and z-scored within block relative to the −1000 to −100 ms pre-cue baseline. Complementary frequency-band analyses applied the same RSA pipeline to theta power (4–8 Hz; 3–5 wavelet cycles) and beta power (17–25 Hz; 6–10 wavelet cycles), and were treated as control analyses to establish the specificity of sigma power in driving the alignment.

Sensor-level spatiotemporal searchlight RSA was performed across the 0 - 4 s post-cue interval using 500-ms sliding windows advanced in 10-ms steps. For each of the 58 EEG sensors, the searchlight comprised the centre sensor and its four nearest neighbouring sensors, defined from three-dimensional electrode coordinates. Within each searchlight and time window, sigma power was averaged across time and represented as trial-level sensor-by-frequency feature vectors. For each pair of trait cues, neural similarity was estimated using valid cross-block trial pairs only. Spearman correlations were computed between trial-level feature vectors for all valid cross-block trial pairs belonging to the two cue items. These correlations were Fisher-z transformed, averaged within each item pair, back-transformed to correlation coefficients, and converted to neural dissimilarity as 1 − r, yielding a local neural RDM for each searchlight and time window.

Self-evaluative RDMs were constructed using the same item order as the neural RDMs, using graded post-sleep item-level self-descriptiveness ratings averaged across the post-sleep immediate and delayed tests. Behavioural dissimilarity between each pair of items was defined as the absolute difference in these averaged ratings, producing a distance-based model of post-sleep self-evaluative structure. Binary endorsement RDMs were used as complementary behavioural models because it indexes categorical endorsement rather than graded or continuous self-evaluation.

At each searchlight and time window, neural-evaluative representational alignment was computed as the Spearman correlation between the vectorised upper triangles of the neural and self-evaluative RDMs. RSA coefficients were Fisher-z transformed before group-level statistical analysis. Back-transformed or raw-scale values were used only for visualisation.

### Spindle detection and probability

Spindle probability was analysed as a sleep event analysis to supplement the sigma-band RSA result, linking the sigma representational finding to a canonical sleep oscillation implicating memory reactivation. Spindles were detected from artifact-free slow-wave N3 sleep EEG during TMR using YASA in the 11–16 Hz range, with event durations constrained to 0.5–3.0 s. We used a correlation-based criterion requiring a minimum sigma-envelope correlation of 0.5 and an RMS threshold of 1.5; no relative-power threshold was applied. Detection was performed over midline spindle channels (Fz, FCz, Cz), baseline-corrected relative to the −1 to 0 s pre-cue interval, and analysed over the 0–4 s post-cue window. Binary event-presence time series were generated for each trial and time point. Spindle probability was computed as the mean binary event-presence value across trials.

### Statistical analysis

Behavioural analyses were performed in R (version 4.2.1). EEG preprocessing and signal analyses were conducted in MATLAB using FieldTrip, EEGLAB and custom scripts. Across analyses, depressive symptoms were modelled primarily as a continuous variable using Beck Depression Inventory-II (BDI-II) scores. Bayesian models are reported using posterior medians, 95% credible intervals (CrIs) and probability of direction (*pd*). We treated pd as a descriptive index of directional certainty rather than as a stand-alone decision criterion (Makowski et al., 2019). For clarity, we refer to pd ≥ 99% as very strong directional evidence, 97.5% ≤ pd < 99% as strong directional evidence, 95% ≤ pd < 97.5% as limited directional evidence, and pd < 95% as weak or inconclusive evidence. For frequentist EEG analyses, statistical inference relied on non-parametric permutation procedures, with cluster-based correction where appropriate.

### Behavioral analysis

The primary behavioural outcomes were trial-wise endorsement positive traits after sleep. To test whether the effect of cueing varied with depressive symptom severity and across time, we fitted Bayesian multilevel Bernoulli regression models to single-trial endorsement responses. Fixed effects included TMR condition (cued versus uncued), Time (Post-sleep versus Delay), continuous BDI-II score, and all interactions with pre-sleep endorsement of the corresponding item included as a baseline covariate and Age and Gender included as participant-level covariates. The primary reported models used participant-level random intercepts and uncorrelated random slopes for TMR and Time, (1+TMR+Time∣∣participant). Alternative random-effects structures were compared using PSIS-LOO and sampler diagnostics; full comparisons are reported in the Supplementary Note 8. Bayesian models were fitted using the brms package interfacing with Stan (Bürkner, 2017). Models were estimated with four chains of 4,000 iterations each, including 1,000 warm-up iterations per chain. Default priors were applied. Convergence was assessed using R^ values and visual inspection of trace plots. For interpretability, conditional TMR effects were also estimated at low (−1 s.d.), mean and high (+1 s.d.) levels of BDI-II and are reported on the response scale as percentage-point differences between cued and uncued traits for the endorsement binary model. All reported Bayesian models converged satisfactorily, with four well-mixed chains, R^ values below 1.01, and adequate bulk and tail effective sample sizes, and no divergent transitions. Posterior predictive checks suggested no major global misfit (Supplementary Fig. 3).

In formula form, the models were:

Endorsement ∼ 1 + baseline endorsement + TMR × Time × BDI-II + Age + Gender + (1 + TMR + Time || participant)

### EEG data statistical analysis

Statistical inference on EEG measures followed two complementary frameworks, selected according to the structure of the dependent variable. For analyses involving continuous time, frequency or sensor dimensions—including cue-elicited spectral power contrasts (Trait − Control), time-resolved raw EEG representational similarity (WI − BI), searchlight neural– behavioural RSA and cue-elicited spindle probability—non-parametric cluster-based permutation tests were used to control for multiple comparisons while preserving the spatiotemporal dependence structure of the data. For item-level behavioural prediction, where the outcome was a discrete endorsement decision, Bayesian mixed-effects regression models were employed.

### Cluster-based permutation tests

Cluster-based permutation tests followed standard non-parametric procedures (Maris & Oostenveld, 2007; Oostenveld et al., 2011). For one-sample tests (e.g., testing group-level time-resolved WI − BI against zero, or cue-elicited spindle-probability changes relative to baseline), sign-flip permutations were used. For paired comparisons (e.g., cue versus control RSA), paired permutation tests were applied. The search space and cluster neighbourhood were defined by the structure of each analysis: channel × frequency × time for cue-elicited spectral power, time for univariate RSA and spindle probability, and sensor × time for searchlight RSA. Cluster-forming thresholds were set at *P* < 0.05, and corrected cluster-level significance was assessed against the permutation-derived null distribution of the maximum cluster statistic. Unless otherwise specified, 2,000 permutations were used.

### Participant-level BDI associations with group-level neural clusters or windows

For significant group-level clusters or windows defined without using BDI-II score, we extracted one mean neural value per participant and tested whether this value was associated with depressive symptom severity. These scalar follow-up analyses used covariate-adjusted permutation regression:

neural value ∼ BDI_z + age_z + gender

The permutation test was performed at the participant level by permuting the residualized BDI-II regressor after accounting for age and gender. These analyses were not cluster-permutation tests, because the continuous search space had already been reduced to one mean value per participant. For spectral power, these values were extracted from the early and late Trait − Control power clusters. For item-specific EEG similarity, the participant-level value was the mean within-minus-between similarity index. For spindle probability, these participant-level BDI association analyses used the baseline-corrected spindle response extracted from the significant group-level spindle response window. We report the BDI regression coefficient, *t*-statistic, partial correlation, confidence interval and participant-level permutation *P* value.

### Bayesian mixed-effects models for item-level behavioural prediction

Item-level behavioural relevance of EEG measures was tested using Bayesian mixed-effects logistic regression models predicting post-sleep endorsement of positive traits. Endorsement was modelled as a binary outcome using a Bernoulli likelihood with a logit link. For each neural measure, the model included the item-level neural predictor, standardized BDI-II score and their interaction, while adjusting for baseline endorsement, post-sleep session, age and gender:

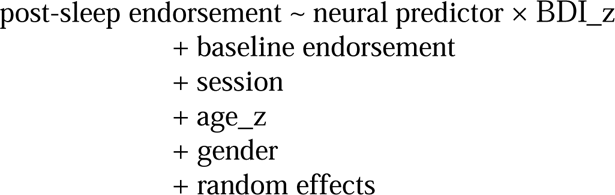

Models included participant-level random effects and item-within-participant random intercepts: (1 + session || participant) + (1 | participant:item)

Spectral power predictors were extracted from the early and late Trait minus Control power clusters, and item-specific EEG predictors were defined as the item-level within-minus-between similarity index. Spindle-based behavioural prediction used two related but distinct predictors. The first predictor was baseline-corrected post-cue spindle probability extracted from the significant spindle cluster, which tested whether stronger cue-elicited spindle responses were associated with post-sleep self-evaluation. The second predictor was raw post-cue spindle probability extracted from the 1.99-2.92 s sigma-band neural-evaluative alignment window. In this alignment-window model, raw pre-cue spindle probability was included as a covariate to test whether post-cue spindle occurrence predicted endorsement beyond baseline spindle propensity. Continuous neural predictors were standardized before modelling. Posterior effects are reported as posterior means, 95% credible intervals and probability of direction.

## Data availability

The de-identified behavioural data and processed EEG-derived measures used for the main analyses are available via Open Science Framework at: https://osf.io/e9nms/overview?view_only=60db934b4ef1498d8b28c9bf537aa106. The full list of personality traits and control words is provided in Supplementary Table 1 and is also included in the repository.

## Code availability

Code used for behavioural and EEG data analyses is available via Open Science Framework at: https://osf.io/e9nms/overview?view_only=60db934b4ef1498d8b28c9bf537aa106.

## Acknowledgements

We thank all participants for their participation. We thank Xingyi Jin, Man Hei Eunice Choi, Pei Yi Tu who assisted with participant recruitment, overnight sleep monitoring, behavioural testing and data quality control.

## Author contributions

Z.Y., Y.M. and X.H. conceived the study. Z.Y., and X.H. designed the experiment. Z.Y., J.W., X.Z., and C.X.C. contributed to data collection, sleep monitoring and EEG procedures. Z.Y., D.C., J.L., J.W., T.X., and X.H. contributed to EEG preprocessing, time-frequency analyses, representational similarity analyses and spindle-event analyses. Z.Y. performed the statistical analyses with input from D.C., J.L., J.W., T.X., P.Q., Y.M. and R.N.Y.C., S.X.L., T.Y., Z.Z., T.M.C.L., Y.M. and X.H.. Z.Y., Y.M. and X.H. wrote the original manuscript. All authors contributed to writing-revision, editing, and approved the final version. X.H. supervised the project and acquired the funding.

