## Supplementary Information for "Sleep reactivation of positive self-evaluation reveals neural-evaluative alignment that varies with depressive symptoms"

### **Supplementary Notes 1-8**

#### **Supplementary Note 1. Cueing effects on continuous self-descriptiveness ratings of positive traits**

For self-descriptiveness ratings of positive traits, we found a significant cueing benefits: median = 0.816, 95% CrI [0.186, 1.436], pd = 99.4% at averaged depressive symptoms when averaging across time. Cueing benefits more clearly expressed at delayed than immediately after TMR: at mean depressive symptoms, the estimated cue benefit was 1.069 rating units at delay (95% CrI [0.225, 1.928], pd = 99.4%), and was 0.564 rating units in the post-sleep session (95% CrI [−0.290, 1.368], pd = 90.2%). Neither the cueing × depression interaction, nor the cueing × time interaction was evident.

To clarify why cueing effects diverged across endorsement and descriptiveness outcomes, we conducted supplementary boundary-based mechanism analyses using the baseline relationship between descriptiveness ratings and endorsement. First, baseline endorsement was regressed on baseline descriptiveness to estimate an empirical endorsement boundary on the descriptiveness scale, defined as the value at which the model-predicted probability of endorsement was .50. Baseline non-endorsed items falling below this boundary were then classified according to their proximity to the boundary: items within 10 scale units below the boundary were treated as near-threshold items, whereas items more than 15 scale units below the boundary were treated as far-below-threshold items. Items falling between these bands were not included in the boundary-zone contrasts.

We then fit three follow-up Bayesian multilevel models. The first model tested endorsement crossing, defined as a transition from non-endorsement at baseline to endorsement at the later test. The second model tested whether the same boundary structure was expressed in descriptiveness ratings, using change in descriptiveness from baseline as the outcome. The third model tested subthreshold strengthening by restricting the rating-change analysis to items that remained below the empirical endorsement boundary at the later test. Each model included cueing condition, standardized BDI score, boundary zone, and their interaction, with standardized age and gender included as covariates. Random intercepts were included for subject–item combinations, and subject-level random slopes for cueing were included where appropriate. Cueing effects were summarized as posterior cue-minus-uncue contrasts at lower and higher depressive symptom severity, defined as BDI = −1 SD and BDI = +1 SD, respectively.

The clearest evidence for endorsement crossing was observed among participants with lower depressive symptom severity for items that were already near the endorsement boundary at baseline. In this condition, cueing increased the probability of endorsement crossing by an estimated 0.393, with the 95% credible interval excluding zero, 95% CrI [0.084, 0.661], pd = 99.1%. The corresponding estimate for lower-BDI participants was smaller for items farther below the boundary, median = 0.123, 95% CrI [−0.035, 0.317], pd = 93.5%. In contrast, endorsement-crossing effects were not clearly expressed among participants with higher depressive symptom severity, either for near-threshold items, median = −0.098, 95% CrI [−0.326, 0.142], pd = 80.1%, or for far-below-threshold items, median = −0.022, 95% CrI [−0.103, 0.056], pd = 72.3%.

The complementary rating-change model was directionally compatible with this boundary-crossing interpretation. Among lower-BDI participants, cue-related increases in descriptiveness were larger for near-threshold items, median = 5.975, 95% CrI [−3.383, 13.695], pd = 91.5%, than for far-below-threshold items, median = 1.089, 95% CrI [−4.377, 6.258], pd = 65.7%. Among higher-BDI participants, the near-threshold estimate was not positive, median = −0.694, 95% CrI [−6.415, 5.091], pd = 60.2%, whereas the far-below-threshold estimate was positive but uncertain, median = 2.121, 95% CrI [−1.453, 5.658], pd = 89.3%.

Finally, the subthreshold rating-increase model examined items that remained below the empirical endorsement boundary at the later test. This analysis provided supportive, but not definitive, evidence that cue-related increases in positive descriptiveness could remain below the endorsement threshold rather than being expressed as endorsement transitions. This pattern was most apparent among higher-BDI participants for far-below-threshold items, median = 2.000, 95% CrI [−0.452, 4.772], pd = 93.6%. The corresponding near-threshold estimate for higher-BDI participants was negative and uncertain, median = −3.123, 95% CrI [−8.795, 2.644], pd = 85.5%. Among lower-BDI participants, the subthreshold model showed a positive but uncertain near-threshold estimate, median = 4.646, 95% CrI [−3.153, 12.582], pd = 86.4%, and little evidence for a far-below-threshold effect, median = −0.528, 95% CrI [−5.006, 3.982], pd = 59.3%.

Taken together, these supplementary analyses support a boundary-based interpretation of the main behavioural dissociation. At lower depressive symptom severity, cue-related effects were most consistent with near-threshold endorsement crossing. At higher depressive symptom severity, cueing effects were less likely to be expressed as categorical endorsement transitions and were more compatible with subthreshold strengthening of positive descriptiveness, particularly for items that began farther below the endorsement boundary. Full posterior summaries are reported in Supplementary Table 7.

#### **Supplementary Note 2. Recall of endorsed traits showed no clear TMR-related change or reliable moderation by depressive symptoms**

Self-evaluative recall was recoded so that a trait counted as remembered only when it had been both endorsed and later recalled; all other traits were coded as forgotten. This conditional definition was used to focus the analysis on self-evaluative memories rather than recall more generally. Trial-level recall was then modeled using a Bayesian hierarchical logistic model including Time, TMR, standardized BDI, and their interactions, with Age and Gender as covariates.

There was no clear evidence that the cueing effect on conditional recall depended reliably on depressive symptom severity, as indicated by the predicted TMR × depression moderation contrast averaged across the post-sleep and delayed tests. The lower-minus-higher BDI contrast was positive but uncertain, median = 0.028, 95% CrI [−0.008, 0.065], pd = 93.9%. Averaging across the post-sleep and delayed tests, cueing produced a small and uncertain increase in conditional recall at lower depressive symptom severity, +2.0 percentage points, 95% CrI [−0.8, 5.0], pd = 92.0%, whereas the effect was close to zero at mean depressive symptom severity, +0.3 percentage points, 95% CrI [−1.3, 2.2], pd = 65.2%, and slightly negative but uncertain at higher depressive symptom severity, −0.8 percentage points, 95% CrI [−2.9, 1.2], pd = 77.3% in the negative direction. The empirical marginal TMR effect, averaged equally across the post-sleep and delayed sessions, was likewise close to zero, +0.7 percentage points, 95% CrI [−1.6, 3.0], pd = 72.1%. Thus, the data provide little support for a reliable TMR effect on recall of endorsed traits or for robust moderation of that effect by depressive symptoms.

#### **Supplementary Note 3. Importance ratings showed no clear TMR-related change or reliable moderation by depressive symptoms**

To examine whether sleep reactivation altered the perceived importance of traits to self-identity, we modeled trial-level importance ratings using a Bayesian Student-*t* mixed model including Time, TMR, standardized BDI, and their interactions, with Age and Gender as covariates.

There was little evidence that the cueing effect on importance ratings depended reliably on depressive symptom severity, as indicated by the predicted TMR × depression moderation contrast averaged across the post-sleep and delayed tests. The lower-minus-higher BDI contrast was close to zero and highly uncertain, median = 0.091 rating units, 95% CrI [−1.555, 1.826], pd = 54.1%. Averaging across the post-sleep and delayed tests, cueing effects on importance ratings were positive but uncertain at lower depressive symptom severity, median = 0.414 rating units, 95% CrI [−0.718, 1.603], pd = 75.5%, at mean depressive symptom severity, median = 0.358 rating units, 95% CrI [−0.541, 1.149], pd = 80.0%, and at higher depressive symptom severity, median = 0.311 rating units, 95% CrI [−0.945, 1.473], pd = 69.3%. The empirical marginal TMR effect, averaged equally across the post-sleep and delayed sessions, was also uncertain, median = 0.358 rating units, 95% CrI [−0.541, 1.149], pd = 80.0%. Overall, these results do not support a reliable TMR effect on importance ratings or robust moderation of that effect by depressive symptom severity.

#### **Supplementary Note 4. No differences in baseline endorsement between cue and uncue positive trait words**

To confirm successful assignment of positive trait words to the cue and uncue conditions, we compared baseline endorsement, descriptiveness, recall, and importance between the two positive-trait sets using participant-level paired tests. There was no evidence of baseline differences on any measure (all *Ps* ≥ 0.153), and all confidence intervals included zero. These results indicate that cue and uncue positive trait words were well matched before sleep. Full statistics are reported in Supplementary Table S5.

#### **Supplementary Note 5. Neural cue responses, depressive symptoms and behavioural updating**

To keep the neural analyses comparable across spectral power, item-specific EEG similarity and spindle probability, we used a common analysis hierarchy. First, we identified group-level neural responses to auditory cueing. For EEG power, this was the Trait − Control time-frequency cluster analysis. For item-specific EEG similarity, this was the within-minus-between multichannel EEG similarity analysis. For spindle probability, this was the cue-locked baseline-corrected spindle probability analysis. These clusters or windows were defined independently of BDI-II score.

Second, for each significant group-level cluster or window, we extracted one mean neural value per participant and tested whether this participant-level value was associated with depressive symptom severity. These analyses used covariate-adjusted permutation regressions:

neural value ~ BDI_z + age_z + gender

The permutation test was performed at the participant level by permuting the residualized BDI-II regressor after accounting for age and gender. These were scalar follow-up tests, not cluster-permutation tests, because each participant contributed one mean value from a previously identified group-level cluster or window.

Across neural measures, BDI-II score did not reliably predict the magnitude of the group-level neural responses. This was true for early Trait − Control power, *b* = 0.004, *t*(58) = 0.31, partial *r* = 0.040, permutation *P* = 0.760; late Trait − Control power, *b* = 0.003, *t*(58) = 0.33, partial *r* = 0.043, permutation *P* = 0.739; item-specific EEG similarity, *b* < 0.001, *t*(58) = 0.04, partial *r* = 0.006, permutation *P* = 0.966; cluster-window spindle probability, *b* = 0.001, *t*(58) = 0.88, partial *r* = 0.114, permutation *P* = 0.370; and fixed-window spindle probability, *b* = 0.002, *t*(58) = 0.67, partial *r* = 0.087, permutation *P* = 0.494. All FDR-adjusted permutation *P*s were ≥ 0.950.

Third, we tested whether item-level neural values predicted post-sleep endorsement using Bayesian mixed-effects models. Each model predicted binary post-sleep endorsement from the neural predictor, standardized BDI-II score and their interaction, while controlling for baseline endorsement, post-sleep session, age and gender, with random effects for participants and participant–item pairs. Early power, late power, item-specific EEG similarity and cluster-window spindle probability did not provide reliable evidence for neural predictor × BDI-II moderation. By contrast, the RSA-aligned fixed spindle window showed evidence for symptom-dependent behavioural relevance: the spindle probability × BDI-II interaction was negative, posterior mean = −0.292, 95% CrI [−0.583, −0.015], pd = 98.0%. Thus, depressive symptoms were not associated with the overall magnitude of cue-elicited neural responses, but moderated how spindle probability related to subsequent positive self-evaluation.

#### **Supplementary Note 6. No BDI-II association with theta- or beta-band RSA**

To test whether the depressive symptom severity-related RSA effect extended beyond the sigma range, we repeated the sensor-level spatiotemporal RSA procedure for theta- and beta-band power. Theta power was defined as 4–8 Hz and estimated using Morlet wavelets with 3–5 cycles across frequencies. Beta power was defined as 17–25 Hz and estimated using 6–10 cycles. Both analyses used the same descriptiveness-rating RDM, searchlight definition, 0–4 s post-cue interval, 500-ms sliding windows, covariates and cluster-based permutation procedure as the primary sigma-band RSA.

Neither theta- nor beta-band RSA showed a cluster-corrected association between BDI-II scores and neural–behavioural representational alignment. These supplementary analyses therefore did not identify additional depressive symptom severity-related representational alignment effects outside the sigma band.

#### **Supplementary Note 7. Midline-overlap follow-up of sigma-band neural–evaluation alignment**

Because the primary RSA effect emerged in the 11–16 Hz sigma range, we conducted a descriptive follow-up using Fz-, FCz- and Cz-centred searchlights that overlapped the primary 47-sensor searchlight cluster. This 3-sensor midline-overlap ROI was chosen to test whether the depression-related neural–evaluation alignment was preserved over frontocentral sensors relevant to NREM sigma/spindle activity.

The midline-overlap ROI reproduced the negative association between depressive symptom severity and neural–evaluation alignment, β = −0.0300, *t* = −3.54, *P* < 0.001. A time-resolved follow-up identified a cluster-corrected negative BDI-II–RSA association from 2.20 to 2.57 s after cue onset, *P* = 0.037. Descriptive lower- and higher-symptom groups showed the same pattern, with stronger alignment in the lower-symptom group, Welch *t*(59.45) = 4.15, *P* < 0.001. These group-based summaries were used only to describe the continuous BDI-II association.

Positive-alignment windows within the lower-symptom group were similar when RSA values were extracted from the full primary 47-sensor cluster and from the restricted midline-overlap ROI. The positive-alignment window extended from 2.13 to 2.81 s in the primary 47-sensor cluster and from 2.22 to 2.59 s in the 3-sensor midline-overlap ROI. These follow-ups indicate that the primary sigma-band RSA effect was preserved in frontocentral sensors relevant to NREM spindle physiology.

#### **Supplementary Note 8. Random-effects model comparison supported the use of a common uncorrelated additive structure across analyses**

We compared four candidate random-effects structures for each behavioural outcome: correlated interaction slopes, uncorrelated interaction slopes, correlated additive slopes, and uncorrelated additive slopes. All candidate structures showed satisfactory sampling behaviour, with no divergent transitions and R^Λ^ values close to 1 throughout (maximum R^Λ^=1.006).

Because the same structure was not uniformly optimal for every outcome, we selected a common parsimonious specification for the main analyses: (1 + TMR + Time || participant). This structure retained participant-specific variability in the two design factors of interest while avoiding the additional complexity of estimating random-effect correlations and higher-order random slopes. It also showed stable convergence and good effective sample size across outcomes. We therefore adopted this common uncorrelated additive structure to maximize stability and comparability across endorsement, descriptiveness, conditional recall, and importance models. Full model-comparison diagnostics are reported in Supplementary Table 6.

### **Supplementary Figures 1-3**

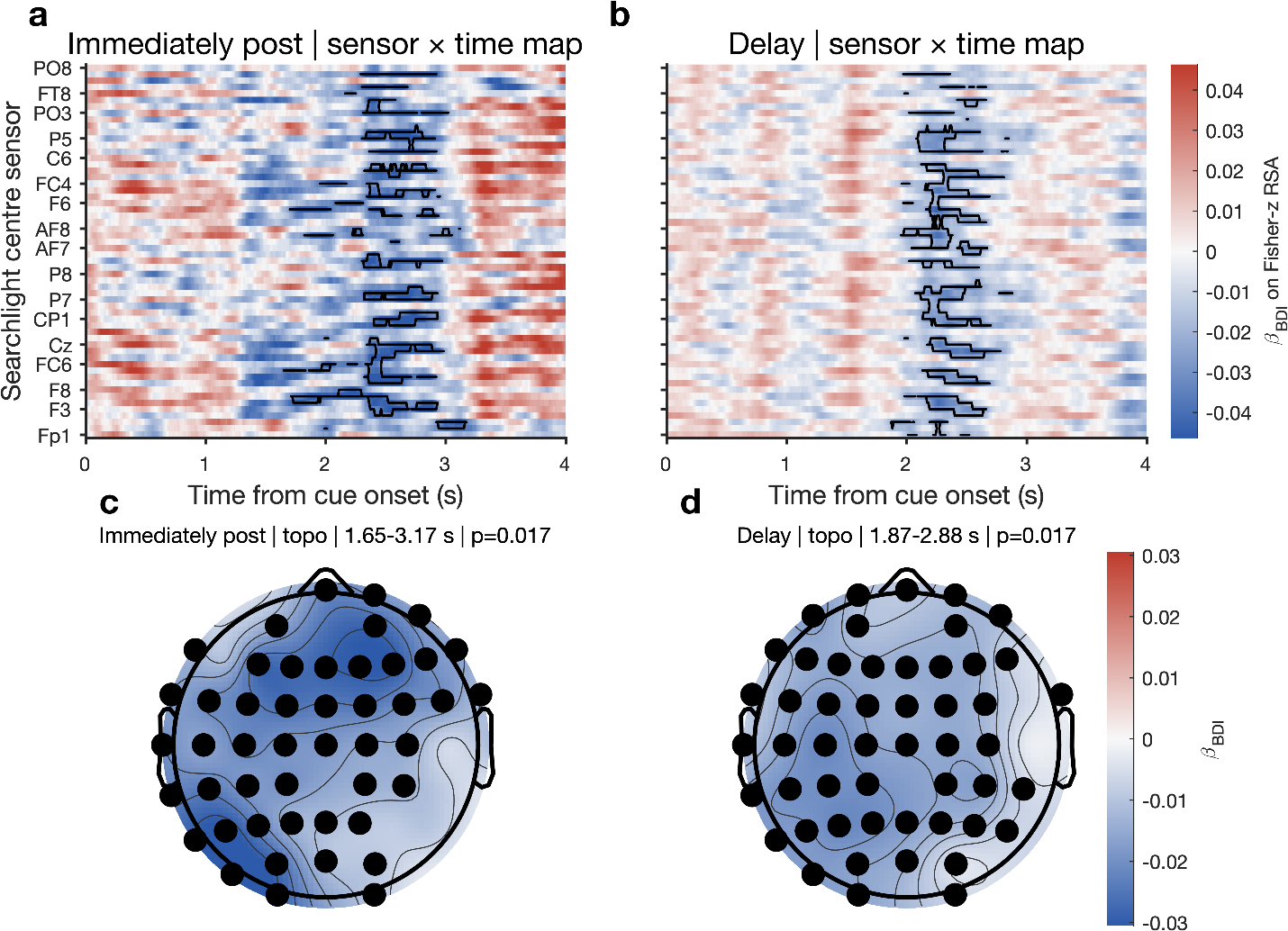

#### **Supplementary Fig. 1 Sigma-band searchlight RSA using immediate and delayed self descriptiveness ratings.**

**a,b,** Sensor × time maps showing the BDI-II regression coefficient for sigma-band neural–behaviour representational alignment using behavioural RDMs derived from raw descriptiveness ratings at the immediate post-TMR assessment (**a**) and delayed assessment (**b**). Colours indicate the BDI-II regression coefficient after adjustment for age and sex/gender. Black contours mark cluster-corrected significant sensor × time clusters.

**c,d,** Topographic distributions of the BDI-II regression coefficient averaged across the corresponding significant cluster windows for the immediate post-TMR (**c**) and delayed (**d**) assessments. Black markers indicate searchlight centre sensors contributing to the significant cluster. Together, these analyses show that the negative depression-related association in sigma-band neural–behaviour alignment was present when behavioural RDMs were constructed separately from immediate and delayed descriptiveness ratings.

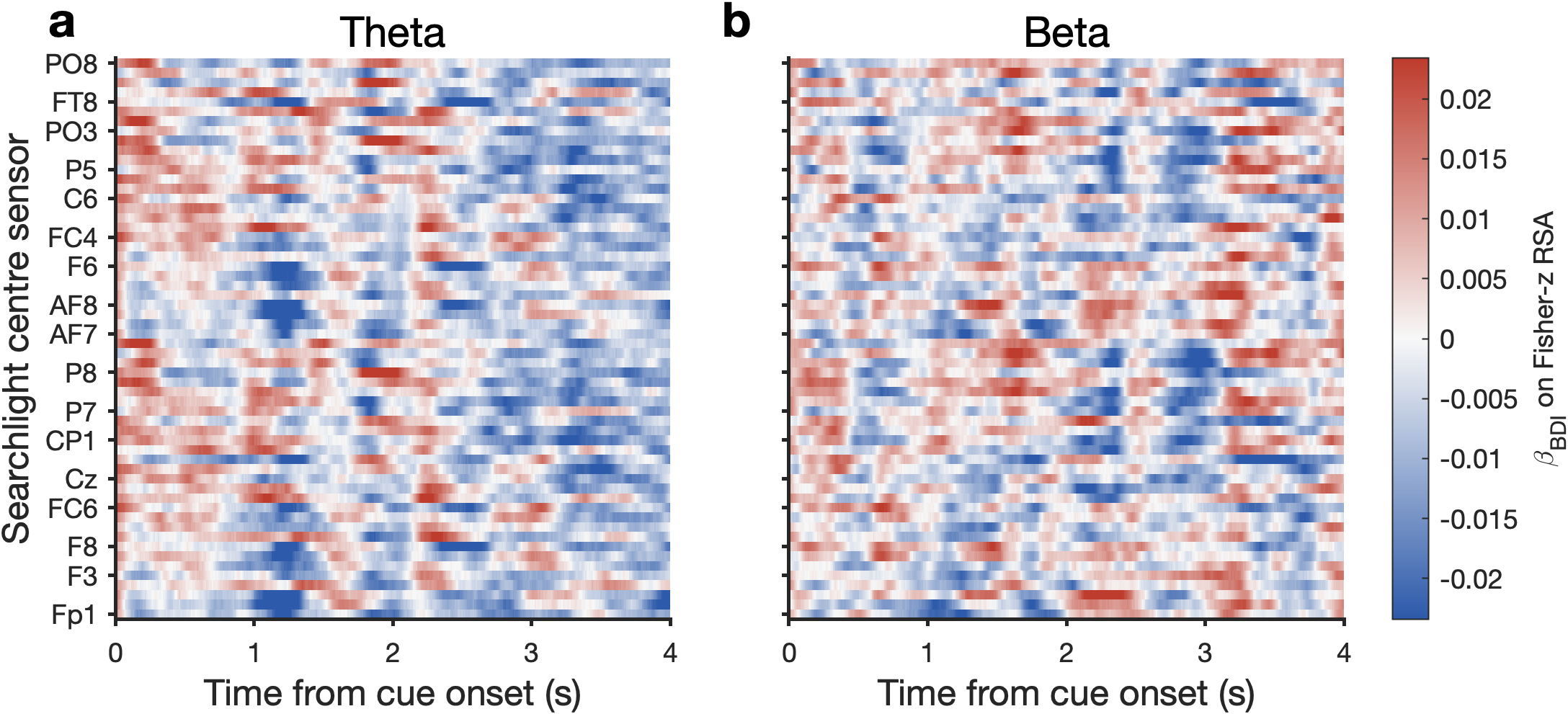

#### **Supplementary Fig. 2 Theta- and beta-band control searchlight RSA.**

**a,b,** Sensor × time maps showing the BDI-II regression coefficient for neural–behaviour representational alignment in the theta band (**a**) and beta band (**b**). The same searchlight RSA procedure as in the sigma-band analysis was applied, with neural RDMs constructed from cue-locked EEG power and correlated with the behavioural RDM. Colours indicate the regression coefficient for BDI-II after adjustment for age and sex/gender, with blue denoting more negative BDI-related associations and red denoting more positive associations. No cluster-corrected depression-related alignment effects were observed in either frequency band.

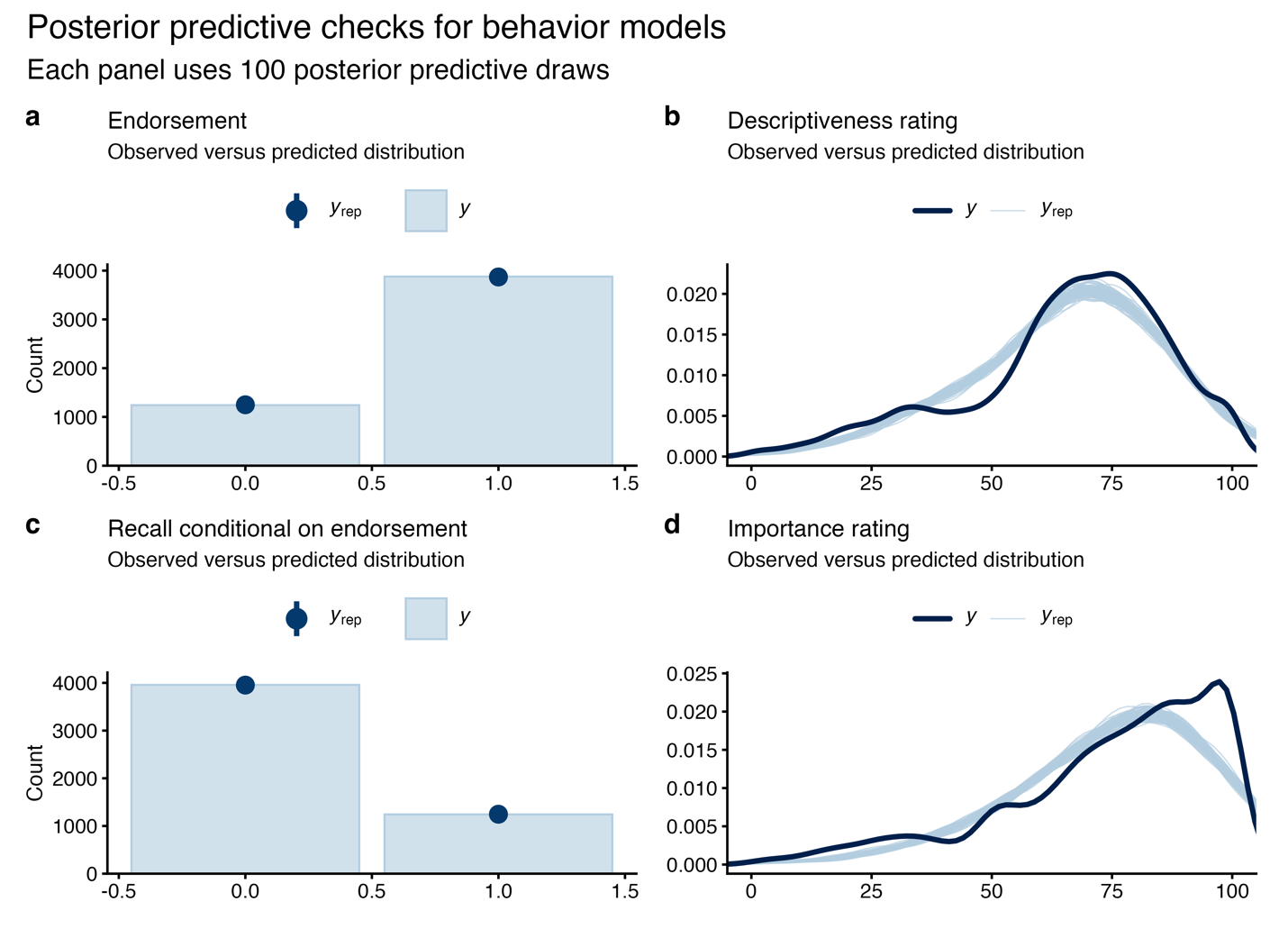

#### **Supplementary Fig. 3 Posterior predictive checks for the behavioural models.**

**a,** Endorsement. **b,** Descriptiveness ratings. **c,** self-evaluative recall. **d,** Importance ratings. Observed distributions (**y**) are compared with posterior predictive replicated data (**yrep**) from the fitted models. In **a** and **c**, light blue bars indicate the observed distributions and dark blue points with intervals indicate replicated counts. In **b** and **d**, the dark blue line indicates the observed distribution, and the light blue lines indicate replicated distributions. The close overlap between observed and replicated data suggests adequate model fit.

### **Supplementary Tables 1-7**

#### **Supplementary Table 1.** word stimuli used in the SRET task

*Note.* Stimuli were categorized as positive stimuli, negative stimuli, and control stimuli.

| **Traditional Chinese** | **Simplified Chinese** | **English translation** | **Valence** |
| --- | --- | --- | --- |
| 樂觀 | 乐观 | optimistic | Positive |
| 成熟 | 成熟 | mature | Positive |
| 體貼 | 体贴 | considerate | Positive |
| 勇敢 | 勇敢 | courageous | Positive |
| 勤奮 | 勤奋 | diligent | Positive |
| 博學 | 博学 | educated | Positive |
| 友善 | 友善 | friendly | Positive |
| 可靠 | 可靠 | reliable | Positive |
| 和藹 | 和蔼 | gracious | Positive |
| 善良 | 善良 | kind | Positive |
| 堅持 | 坚持 | persistent | Positive |
| 外向 | 外向 | outgoing | Positive |
| 好學 | 好学 | studious | Positive |
| 寬容 | 宽容 | tolerant | Positive |
| 忠誠 | 忠诚 | loyal | Positive |
| 快樂 | 快乐 | happy | Positive |
| 慷慨 | 慷慨 | generous | Positive |
| 無私 | 无私 | unselfish | Positive |
| 有趣 | 有趣 | interesting | Positive |
| 單純 | 单纯 | innocent | Positive |
| 正直 | 正直 | righteous | Positive |
| 活潑 | 活泼 | vivacious | Positive |
| 溫暖 | 温暖 | warm | Positive |
| 熱心 | 热心 | obliging | Positive |
| 熱情 | 热情 | enthusiastic | Positive |
| 獨立 | 独立 | self-reliant | Positive |
| 理智 | 理智 | rational | Positive |
| 直率 | 直率 | direct | Positive |
| 真誠 | 真诚 | sincere | Positive |
| 積極 | 积极 | active | Positive |
| 聰明 | 聪明 | smart | Positive |
| 能幹 | 能干 | capable | Positive |
| 自信 | 自信 | self-confident | Positive |
| 節儉 | 节俭 | thrifty | Positive |
| 誠實 | 诚实 | honest | Positive |
| 謙卑 | 谦卑 | humble | Positive |
| 謙虛 | 谦虚 | modest | Positive |
| 謹慎 | 谨慎 | cautious | Positive |
| 進取 | 进取 | enterprising | Positive |
| 風趣 | 风趣 | amusing | Positive |
| 不滿 | 不满 | resentful | Negative |
| 專橫 | 专横 | domineering | Negative |
| 傲慢 | 傲慢 | insolent | Negative |
| 冒犯 | 冒犯 | offensive | Negative |
| 冷淡 | 冷淡 | cold | Negative |
| 刻薄 | 刻薄 | unkind | Negative |
| 压抑 | 压抑 | hostile | Negative |
| 吝嗇 | 吝啬 | mean | Negative |
| 善忘 | 善忘 | absent-minded | Negative |
| 固執 | 固执 | obstinate | Negative |
| 失職 | 失职 | negligent | Negative |
| 妒忌 | 妒忌 | envious | Negative |
| 孤僻 | 孤僻 | antisocial | Negative |
| 幼稚 | 幼稚 | childish | Negative |
| 惡毒 | 恶毒 | malicious | Negative |
| 悲觀 | 悲观 | pessimistic | Negative |
| 愚蠢 | 愚蠢 | foolish | Negative |
| 懶散 | 懒散 | listless | Negative |
| 懦弱 | 懦弱 | cowardly | Negative |
| 抱怨 | 抱怨 | complaining | Negative |
| 挑剔 | 挑剔 | fault-finding | Negative |
| 無知 | 无知 | ungracious | Negative |
| 易怒 | 易怒 | irritable | Negative |
| 暴躁 | 暴躁 | cruel | Negative |
| 浮躁 | 浮躁 | weak | Negative |
| 焦慮 | 焦虑 | heartless | Negative |
| 狠心 | 狠心 | sly | Negative |
| 狡猾 | 狡猾 | deceitful | Negative |
| 狡詐 | 狡诈 | dominating | Negative |
| 疏忽 | 疏忽 | neglectful | Negative |
| 粗心 | 粗心 | thoughtless | Negative |
| 緊張 | 紧张 | nervous | Negative |
| 膚淺 | 肤浅 | superficial | Negative |
| 膽怯 | 胆怯 | timid | Negative |
| 自私 | 自私 | selfish | Negative |
| 自負 | 自负 | self-conceited | Negative |
| 虛偽 | 虚伪 | insincere | Negative |
| 貪婪 | 贪婪 | greedy | Negative |
| 邋遢 | 邋遢 | untidy | Negative |
| 魯莽 | 鲁莽 | hot-headed | Negative |
| 交互 | 交互 | interactive | Control |
| 先决 | 先决 | prerequisite | Control |
| 初创 | 初创 | start-up | Control |
| 初始 | 初始 | initial | Control |
| 历任 | 历任 | successive | Control |
| 合算 | 合算 | cost-effective | Control |
| 四方 | 四方 | four-sided | Control |
| 大幅 | 大幅 | substantial | Control |
| 广义 | 广义 | broad | Control |
| 细长 | 细长 | slender | Control |

#### **Supplementary** Table 2. Descriptive statistics for questionnaire measures

| **Measure** | **Abbrev** | **M** | **SD** | **Min** | **Max** |
| --- | --- | --- | --- | --- | --- |
| State-Trait Anxiety Inventory: State | STAI-State | 36.34 | 8.20 | 21.00 | 56.00 |
| State-Trait Anxiety Inventory: Trait | STAI-Trait | 42.55 | 10.01 | 25.00 | 64.00 |
| Barratt Impulsiveness Scale: Attentional | BIS-Att | 14.23 | 3.14 | 8.00 | 23.00 |
| Barratt Impulsiveness Scale: Motor | BIS-Motor | 20.69 | 3.72 | 13.00 | 33.00 |
| Barratt Impulsiveness Scale: Non-planning | BIS-Nonplan | 22.91 | 4.95 | 12.00 | 37.00 |
| Barratt Impulsiveness Scale: Total | BIS-Total | 57.83 | 10.35 | 39.00 | 93.00 |
| Emotion Regulation Questionnaire: Reappraisal | ERQ-Reappraisal | 31.60 | 5.81 | 19.00 | 42.00 |
| Emotion Regulation Questionnaire: Suppression | ERQ-Suppression | 14.14 | 5.36 | 4.00 | 27.00 |
| Emotion Regulation Questionnaire: Total | ERQ-Total | 45.74 | 7.90 | 29.00 | 62.00 |
| Five Facet Mindfulness Questionnaire: Observing | FFMQ-Observing | 25.11 | 5.00 | 13.00 | 34.00 |
| Five Facet Mindfulness Questionnaire: Describing | FFMQ-Describing | 27.17 | 5.41 | 16.00 | 37.00 |
| Five Facet Mindfulness Questionnaire: Acting with Awareness | FFMQ-Awareness | 26.08 | 7.07 | 8.00 | 40.00 |
| Five Facet Mindfulness Questionnaire: Nonjudging | FFMQ-Nonjudging | 23.34 | 5.33 | 13.00 | 34.00 |
| Five Facet Mindfulness Questionnaire: Nonreactivity | FFMQ-Nonreactivity | 20.72 | 4.12 | 12.00 | 29.00 |
| Five Facet Mindfulness Questionnaire: Total | FFMQ-Total | 122.42 | 15.65 | 83.00 | 165.00 |
| Life Orientation Test-Revised | LOT-R | 22.94 | 3.50 | 13.00 | 28.00 |
| Narcissistic Personality Inventory | NPI | 7.28 | 1.64 | 4.00 | 11.00 |
| Rosenberg Self-Esteem Scale (raw sum) | RSES | 35.55 | 7.97 | 15.00 | 49.00 |
| Big Five: Extraversion | BF-Extraversion | 24.51 | 6.00 | 11.00 | 36.00 |
| Big Five: Agreeableness | BF-Agreeableness | 33.78 | 4.93 | 25.00 | 43.00 |
| Big Five: Conscientiousness | BF-Conscientiousness | 28.83 | 6.97 | 11.00 | 42.00 |
| Big Five: Neuroticism | BF-Neuroticism | 25.08 | 5.95 | 9.00 | 39.00 |
| Big Five: Openness | BF-Openness | 36.34 | 6.17 | 19.00 | 48.00 |
| Big Five: Total | BF-Total | 148.54 | 12.13 | 119.00 | 173.00 |
| Ruminative Responses Scale | RRS | 48.74 | 11.19 | 28.00 | 79.00 |
| Beck Depression Inventory (0-63 total) | BDI | 9.48 | 7.80 | 0.00 | 26.00 |

#### **Supplementary Table 3.** Descriptive statistics for total sleep duration and sleep-stage duration

| Measure | M | SD | Min | Max |
| --- | --- | --- | --- | --- |
| Total sleep duration (min) | 432.62 | 51.61 | 220.5 | 493.0 |
| Wake (min) | 42.95 | 48.55 | 4.5 | 255.0 |
| N1 (min) | 23.14 | 15.79 | 3.0 | 81.5 |
| N2 (min) | 224.07 | 36.31 | 141.5 | 287.0 |
| N3 (min) | 92.68 | 23.61 | 48.5 | 155.0 |
| REM (min) | 92.74 | 29.92 | 22.0 | 158.0 |

Note. Descriptive statistics are restricted to participants with at least 360 min of raw overnight EEG recording duration (N = 53 of 65). The remaining 12 participants were excluded from this table because EEG recordings were lost following TMR or during the latter part of the night, resulting in less than 360 min of recorded data. All values are reported in minutes.

#### **Supplementary Table 4** Overview of primary, follow-up, and supplementary behavioral models

| **Model** | **Analysis tier** | **Outcome** | **Fixed effects** | **Random effects** | **Family** |
| --- | --- | --- | --- | --- | --- |
| M1 | Primary | Endorsement | TMR × Time× BDI_z + baseline_YN + Age_z + Gender | (1 + TMR + Time\|\| SubjectID) | Bernoulli |
| M2 | Follow-up | Descriptiveness rating | TMR × Time× BDI_z + Baseline Descriptiveness + Age_z + Gender | (1 + TMR + Time\|\| SubjectID) | Student-t |
| M3 | Follow-up | Recall | baseline_recall + Age_z + Gender + TMR × Time× BDI_z | (1 + Time+ TMR \|\| SubjectID) | Bernoulli |
| M4 | Follow-up | Importance rating | baseline_importance + Age_z + Gender + TMR × Time× BDI_z | (1 + Time+ TMR \|\| SubjectID) | Student-t |
| M5 | Exploratory decomposition | Endorsement | Age_z + Gender + Time× Valence × BDI_z | (1 + Time+ Valence \|\| SubjectID) | Bernoulli |
| M6 | Exploratory decomposition | Descriptiveness rating | Age_z + Gender + Time× Valence × BDI_z | (1 + Time+ Valence \|\| SubjectID) | Student-t |
| M7 | Exploratory decomposition | Recall | Age_z + Gender + Time× Valence × BDI_z | (1 + Time+ Valence \|\| SubjectID) | Bernoulli |
| M8 | Exploratory decomposition | Importance rating | Age_z + Gender + Time× Valence × BDI_z | (1 + Time+ Valence \|\| SubjectID) | Student-t |
| M9 | Supplementary mechanism A | Endorsement crossing | TMR × BDI_z × boundary_zone + Age_z + Gender | (1 + TMR \|\| SubjectID) + (1 \| SubItemID) | Bernoulli |
| M10 | Supplementary mechanism B | Rating increase bridge | TMR × BDI_z × boundary_zone + Age_z + Gender | (1 + TMR \|\| SubjectID) + (1 \| SubItemID) | Student-t |
| M11 | Supplementary mechanism C | Below-threshold rating increase | TMR × BDI_z × boundary_zone + Age_z + Gender | (1 + TMR \|\| SubjectID) + (1 \| SubItemID) | Student-t |

*Note.* BDI_z = standardized depressive symptom severity (BDI-II). Time refers to post-TMR and delay in the TMR models and to baseline, post-TMR, and delay in the valence-decomposition models, as applicable. SubItemID = interaction(SubjectID, ItemID). In the supplementary mechanism analyses, the endorsement threshold was estimated from the baseline relationship between descriptiveness ratings and endorsement. primary_mech_dat included items below this threshold at baseline, not endorsed at baseline, and classified as either near-threshold or far-below-threshold. primary_subthreshold_dat was further restricted to items that remained below threshold at the later test.

#### **Supplementary Table 5** Baseline equivalence between cue and uncue positive trait words

| Outcome | Cue M | Uncue M | Mean difference | t(64) | p | 95% CI | dz |
| --- | --- | --- | --- | --- | --- | --- | --- |
| Endorsement | 0.749 | 0.737 | 0.012 | 0.95 | .344 | [-0.013, 0.036] | 0.118 |
| Descriptiveness rating | 65.515 | 66.045 | -0.531 | -1.45 | .153 | [-1.264, 0.203] | -0.179 |
| Recall | 0.313 | 0.311 | 0.002 | 0.49 | .626 | [-0.007, 0.012] | 0.061 |
| Importance rating | 73.756 | 73.577 | 0.179 | 0.25 | .807 | [-1.276, 1.634] | 0.030 |

*Note.* Cue and uncue sets were compared within participants using paired-samples t tests restricted to positive traits at baseline. Mean difference = cue minus uncue. CI = confidence interval. N = 65 participants.

#### **Supplementary Table 6** Comparison of candidate random-effects structures across reported behavioural outcomes

**Panel A. Predictive comparison of candidate random-effects structures**

| **Outcome** | **Structure** | **Formula** | **LOOIC** | **ΔLOOIC** | **Stacking weight** |
| --- | --- | --- | --- | --- | --- |
| Endorsement | Correlated interaction | (1 + Time * TMR \| SubjectID) | 3462.9 | 1.3 | 0.000 |
|  | Uncorrelated interaction | (1 + Time * TMR \|\| SubjectID) | 3461.7 | 0.0 | 0.710 |
|  | Correlated additive | (1 + Time + TMR \| SubjectID) | 3465.5 | 3.8 | 0.000 |
|  | **Uncorrelated additive†** | **(1 + Time + TMR \|\| SubjectID)** | **3463.3** | **1.6** | **0.290** |
| Descriptiveness ratings | Correlated interaction | (1 + Time * TMR \| SubjectID) | 40257.4 | 5.4 | 0.000 |
|  | Uncorrelated interaction | (1 + Time * TMR \|\| SubjectID) | 40254.0 | 2.0 | 0.001 |
|  | Correlated additive | (1 + Time + TMR \| SubjectID) | 40255.4 | 3.5 | 0.001 |
|  | **Uncorrelated additive†** | **(1 + Time + TMR \|\| SubjectID)** | **40252.0** | **0.0** | **0.999** |
| Conditional recall | Correlated interaction | (1 + Time * TMR \| SubjectID) | 4982.9 | 0.0 | 0.697 |
|  | Uncorrelated interaction | (1 + Time * TMR \|\| SubjectID) | 4985.9 | 3.1 | 0.000 |
|  | Correlated additive | (1 + Time + TMR \| SubjectID) | 4983.5 | 0.7 | 0.301 |
|  | **Uncorrelated additive†** | **(1 + Time + TMR \|\| SubjectID)** | **4984.6** | **1.7** | **0.002** |
| Importance ratings | Correlated interaction | (1 + Time * TMR \| SubjectID) | 42587.0 | 4.5 | 0.000 |
|  | Uncorrelated interaction | (1 + Time * TMR \|\| SubjectID) | 42583.5 | 1.1 | 0.208 |
|  | Correlated additive | (1 + Time + TMR \| SubjectID) | 42587.4 | 5.0 | 0.000 |
|  | **Uncorrelated additive†** | **(1 + Time + TMR \|\| SubjectID)** | **42582.4** | **0.0** | **0.792** |

**Panel B. Sampling diagnostics for candidate random-effects structures**

| **Outcome** | **Structure** | **Max R̂** | **Min bulk ESS** | **Min tail ESS** | **Divergences** |
| --- | --- | --- | --- | --- | --- |
| Endorsement | Correlated interaction | 1.006 | 1,929 | 3,369 | 0 |
|  | Uncorrelated interaction | 1.002 | 1,888 | 3,112 | 0 |
|  | Correlated additive | 1.003 | 1,624 | 2,369 | 0 |
|  | **Uncorrelated additive†** | **1.003** | **2,073** | **2,328** | **0** |
| Descriptiveness ratings | Correlated interaction | 1.001 | 2,835 | 4,662 | 0 |
|  | Uncorrelated interaction | 1.002 | 2,499 | 4,487 | 0 |
|  | Correlated additive | 1.002 | 2,666 | 4,485 | 0 |
|  | **Uncorrelated additive†** | **1.002** | **2,957** | **5,058** | **0** |
| Conditional recall | Correlated interaction | 1.001 | 2,397 | 4,306 | 0 |
|  | Uncorrelated interaction | 1.002 | 1,945 | 3,498 | 0 |
|  | Correlated additive | 1.002 | 1,665 | 3,598 | 0 |
|  | **Uncorrelated additive†** | **1.002** | **2,059** | **3,717** | **0** |
| Importance ratings | Correlated interaction | 1.003 | 1,782 | 2,315 | 0 |
|  | Uncorrelated interaction | 1.003 | 2,768 | 3,122 | 0 |
|  | Correlated additive | 1.003 | 1,606 | 1,929 | 0 |
|  | **Uncorrelated additive†** | **1.001** | **2,634** | **2,426** | **0** |

**Panel C. Family comparison for descriptiveness ratings under the common selected random-effects structure**

| **Family** | **LOOIC** | **ΔLOOIC** | **Stacking weight** | **Max R̂** | **Min bulk ESS** | **Min tail ESS** | **Divergences** |
| --- | --- | --- | --- | --- | --- | --- | --- |
| **Student** | **40252.0** | **0.0** | **0.956** | **1.002** | **2,957** | **5,058** | **0** |
| Gaussian | 40882.4 | 630.4 | 0.044 | 1.001 | 3,540 | 6,000 | 0 |

*Note.* LOOIC = leave-one-out information criterion; ΔLOOIC = difference from the lowest LOOIC within outcome or panel; higher stacking weights indicate greater relative predictive support. † Uncorrelated additive (1 + Time + TMR || SubjectID) was adopted as the common random-effects structure for all reported models to preserve comparability across outcomes because its predictive performance was similar to the alternatives. Panel C compares Gaussian and Student families for descriptiveness ratings under this common selected random-effects structure; the Student family was retained for the reported analyses.

#### **Supplementary Table 7** Posterior Summary for Boundary-Based Mechanism Analyses

| **Contrast** | **Estimate** | **95% CrI** | **pd** |
| --- | --- | --- | --- |
| **Endorsement crossing** | | | |
| Low BDI (-1 SD); Near threshold | 0.393 | [0.084, 0.661] | 0.991 |
| Low BDI (-1 SD); Far below threshold | 0.123 | [-0.035, 0.317] | 0.935 |
| High BDI (+1 SD); Near threshold | -0.098 | [-0.326, 0.142] | 0.801 |
| High BDI (+1 SD); Far below threshold | -0.022 | [-0.103, 0.056] | 0.723 |
| **Rating increase support** | | | |
| Low BDI (-1 SD); Near threshold | 5.975 | [-3.383, 13.695] | 0.915 |
| Low BDI (-1 SD); Far below threshold | 1.089 | [-4.377, 6.258] | 0.657 |
| High BDI (+1 SD); Near threshold | -0.694 | [-6.415, 5.091] | 0.602 |
| High BDI (+1 SD); Far below threshold | 2.121 | [-1.453, 5.658] | 0.893 |
| **Subthreshold rating increase** | | | |
| Low BDI (-1 SD); Near threshold | 4.646 | [-3.153, 12.582] | 0.864 |
| Low BDI (-1 SD); Far below threshold | -0.528 | [-5.006, 3.982] | 0.593 |
| High BDI (+1 SD); Near threshold | -3.123 | [-8.795, 2.644] | 0.855 |
| High BDI (+1 SD); Far below threshold | 2.000 | [-0.452, 4.772] | 0.936 |

*Note.* Rows report cue-versus-uncue estimates for the primary probe contrasts used in Supplementary Note 6. Endorsement crossing is expressed on the cue-benefit probability scale; the two rating models are expressed on the raw rating-change scale. Estimate = posterior median; CrI = credible interval; pd = probability of direction.
